# Accurate detection of tumor clonality and ongoing expansion mode from genomic data

**DOI:** 10.64898/2026.06.15.732415

**Authors:** Yanjie Chen, Roman Jaksik, Peter Terranova, Sara El Baghdadi, Andrew Koval, Monika K. Kurpas, Simon Tavaré, Marek Kimmel, Khanh N. Dinh

## Abstract

Recent evidence shows that despite considerable effort, currently available algorithms for estimating intratumor heterogeneity (ITH) remain limited. We developed DECODE (<u>De</u>ciphering <u>C</u>ancer <u>O</u>rigin from <u>D</u>NA <u>E</u>volution), a novel mutation clustering method that incorporates the impact of sample-specific sequencing coverage and mutation calling biases. On synthetic data, DECODE outperformed existing methods across multiple clonality metrics and accurately detected and characterized the neutral tail in the site frequency spectrum (SFS), which encodes the tumor’s ongoing expansion mode. In acute myeloid leukemia, accounting for the neutral tail enabled DECODE to yield more parsimonious clonal decompositions that align more closely with known subclonal dynamics that drive relapse. Applied to data from The Cancer Genome Atlas, DECODE not only detected a neutral SFS tail in most samples across tumor types but also uncovered a clinically meaningful link between ITH and survival in low-grade glioma. By jointly inferring clonality and expansion mode, DECODE provides two complementary and prognostically relevant readouts of tumor evolution from single tumor genomic samples.

## 1 Introduction

Intra-tumor heterogeneity (ITH) has been consistently associated with worse cancer patient outcome [Andor et al., 2016], drug resistance [McGranahan and Swanton, 2017, Dagogo-Jack and Shaw, 2018], and increased risk of relapse and metastasis across different cancer types [Turajlic and Swanton, 2016, Caswell and Swanton, 2017]. Subclonal reconstruction has accordingly become a pillar of cancer evolution research, with growing applications in patient stratification and treatment design [Tarabichi et al., 2021]. Many such methods operate on the site frequency spectrum (SFS), the distribution of somatic mutations by their variant allele frequency (VAF) within a tumor. The shape of the SFS encodes evolutionary history. Clonal mutations form a cluster at high VAFs, and each subclone contributes a lower-VAF cluster. Tumor deconvolution decomposes this spectrum to recover the number and sizes of subclones present. However, despite the proliferation of subclonal reconstruction algorithms, recent studies have exposed substantial limitations that remain unaddressed. The ICGC-TCGA DREAM Somatic Mutation Calling Tumor Heterogeneity and Evolution Challenge found that, among 31 single-sample mutation decomposition algorithms, no single method achieved top performance across all aspects of clonal reconstruction [Salcedo et al., 2025]. Notably, the authors found that algorithm choice influenced performance more strongly than any tumor feature, including purity, copy-number state, and sequencing depth. In addition, ensemble strategies were unable to outperform the best individual methods. Together, these results underline an urgent need for more accurate and robust methods to estimate ITH from cancer genomic data.

We hypothesized that addressing two often-overlooked factors can improve subclonal reconstruction. First, regardless of whether the tumor evolves neutrally or undergoes selective sweeps, neutral mutations that accumulate as cells divide generate a characteristic power-law tail in the SFS [Williams et al., 2016]. The exponent of the SFS tail distribution quantifies the ongoing rate of mutation accumulation relative to cell division, and thus reflects the tumor’s expansion dynamics. Population genetics theory shows that the tail power takes the value 1 in constant-size populations [Watterson, 1975] and 2 under exponential growth [Griffiths and Tavaré, 1998]. To our knowledge, MOBSTER currently remains the only method that explicitly models the neutral tail [Caravagna et al., 2020]. Indeed, the DREAM Challenge found that applying MOBSTER to remove tail mutations before deconvolution reduces the misassignment of neutral variants to spurious subclones [Salcedo et al., 2025]. However, MOBSTER’s authors noted that the algorithm requires high sample purity and coverage of ≥ 100× to detect the tail reliably, exceeding the depth of most available datasets [Caravagna et al., 2020]. The second critical but frequently neglected factor is that sample-specific features can strongly distort the SFS. We previously showed that sequencing coverage, particularly when low or non-uniform as typical of real data, strongly shapes both the dispersion of mutation clusters and the form of the observed SFS tail [Dinh et al., 2020]. Additionally, mutation callers typically discard a large fraction of mutations at low frequencies, which disproportionately depletes and distorts the empirical SFS tail [Krøigård et al., 2016]. Failure to account for these biases may explain why many current methods infer clonality inaccurately and why MOBSTER cannot reliably detect the tail in low-coverage data.

## 2 Results

### 2.1 DECODE incorporates corrections for sequencing coverage and variant calling to improve detection of mutation clusters and neutral tails

We developed DECODE (<u>De</u>ciphering <u>C</u>ancer <u>O</u>rigin from <u>D</u>NA <u>E</u>volution), a novel mutation clustering algorithm based on our previous mathematical results in [Dinh et al., 2020]. DECODE assumes that a sample’s exact site frequency spectrum (SFS) consists of *H* clusters, formed of mutations present in the most recent common ancestors (MRCAs) of each subclone, and a neutral tail containing other mutations that arise as cells of all subclones divide (Fig. 1A). The expected number of mutations present in *k* cells in a sample of *n* cells thus follows

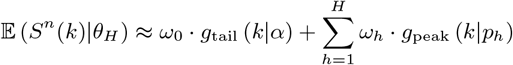

where the *ω*_0_ mutations in the tail follow the power law *g*_tail_ (*k*|*α*) ∝ 1*/k*^*α*^ and the *ω*_*h*_ mutations in each cluster *h* follow the binomial distribution *g*_peak_ (*k*|*p*_*h*_) with mean variant allele frequency (VAF) *p*_*h*_. Given sequencing coverage *φ*, where *φ*(*r*) is the probability that a genomic site is sequenced with a total of *r* reads, we have shown that the expected mutation count with observed VAF in a bin (*f*_1_, *f*_2_] ⊂ [0, 1] follows [Dinh et al., 2020]

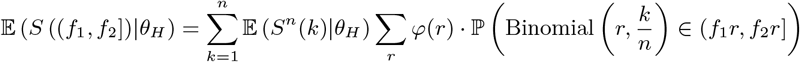

**Figure 1:**
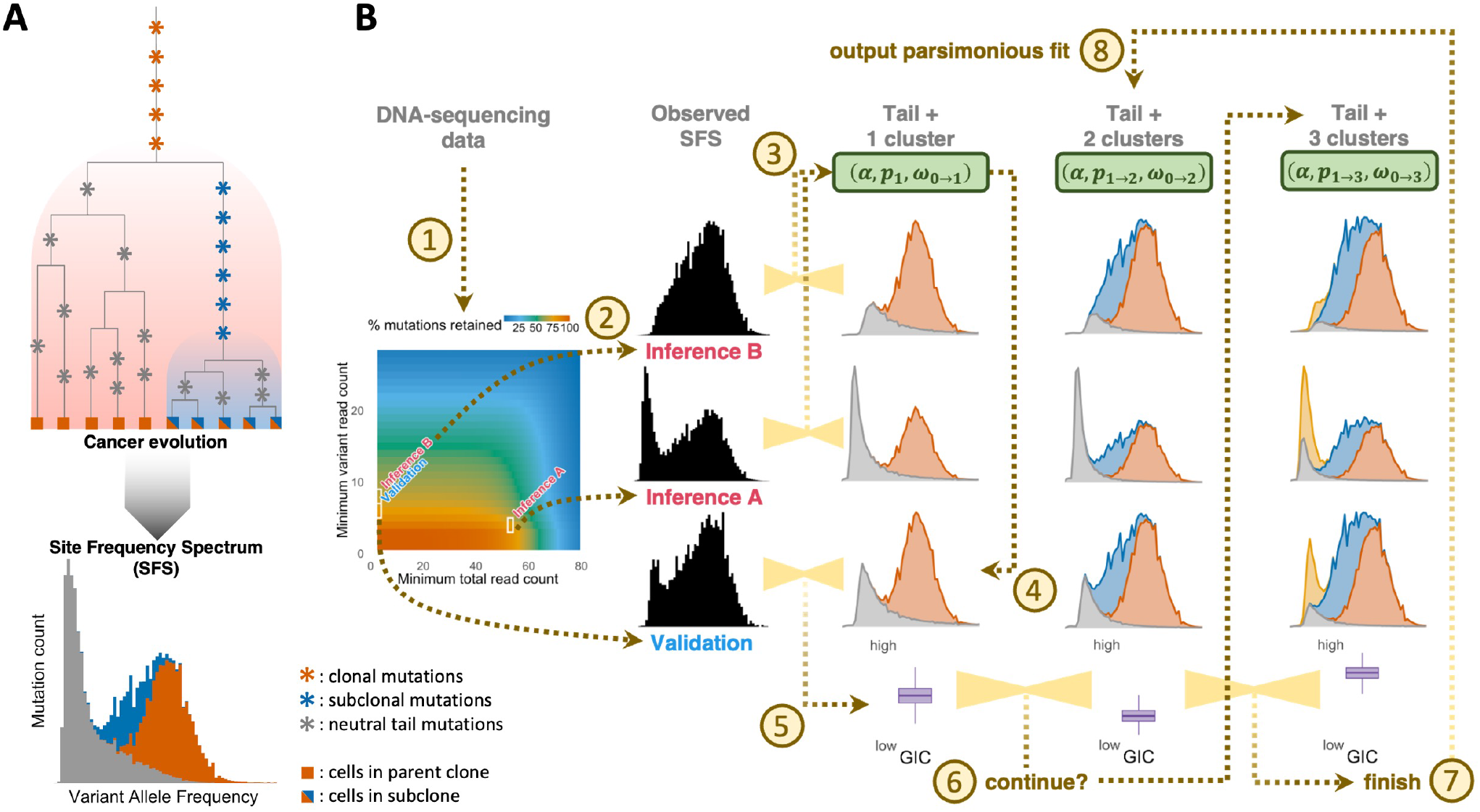
**A**: DECODE’s model for clonal evolution and the corresponding Site Frequency Spectrum (SFS). Clonal mutations are present in a tumor’s MRCA and shared across all cancer cells, forming the truncal cluster in the SFS. Each subclone results from an ongoing selection sweep, where the mutations in the subclone’s MRCA are present in its cells and constitute a subclonal cluster at lower Variant Allele Frequencies (VAFs). Mitoses of cells within all subclones induce additional mutations, resulting in a neutral tail in the lowest-VAF region of the SFS. **B**: Schematic overview of DECODE’s algorithm. For each genomic sample, DECODE extracts the SFS from three filtering strategies, based on variant and total read counts (***step 1-2***). For each cluster count *H*, it estimates model parameters with ABC-SMC-DRF from two inference SFS subsamples (***step 3***), then finds the Generalized Information Criterion (GIC) density based on the predicted SFS for the validation subsample (***step 4-5***). The GIC distributions from successive *H*’s are compared to find the parsimonious fit (***step 6-8***).

The formula can be further modified to accommodate data filtering, which is a critical step in mutation calling and tends to predominantly remove mutations with few variant reads or low coverage in order to reduce the rate of false positives [Krøigård et al., 2016]. Given thresholds (*L, M*) where mutations with fewer than *L* variant reads or *M* total reads are removed, the expected mutation count in bin (*f*_1_, *f*_2_] becomes

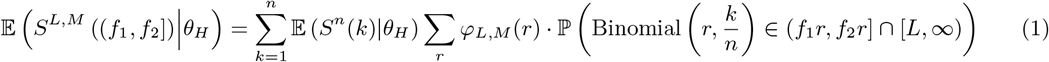

where the coverage distribution *φ*_*L,M*_ (*r*) is computed from the retained mutations. We observed that thresholds *L* and *M* impact the SFS in different ways [Dinh et al., 2020]. Removing mutations with low support (i.e., increasing *L*) mainly reduces the tail, as it is composed of mutations with the lowest VAFs (Fig. 1A). Meanwhile, filtering based on coverage (e.g., by altering *M*) eliminates mutations across the SFS uniformly, and thus only reduces the scale of the SFS without significantly changing its shape.

DECODE takes advantage of our mathematical framework to seek the parsimonious SFS decomposition that is consistent across different data filtering routines. Given a DNA-sequencing sample, it first selects thresholds (*L, M*) for three different data subsamples, termed *inference A, inference B* and *validation* (***step 1***, Fig. 1B). The variant read count threshold is different in each subset, so the SFS from each filtered subsample assumes a different shape (***step 2***, Fig. 1B).

For a given cluster count *H*, DECODE infers the parameters *θ*_*H*_ = (*α, p*_1_, … , *p*_*H*_ , *ω*_0_, *ω*_1_, … , *ω*_*H*_), which characterize the exponent of the tail, each cluster’s mean VAF, and the mutation count in each component. The estimation is obtained via ABC-SMC-DRF (Approximate Bayesian Computation sequential Monte Carlo with distributional random forest) [Dinh et al., 2025], a general likelihood-free inference method that we recently introduced. ABC-SMC-DRF is modified to maintain cluster ordering, where cluster 1 is truncal and cluster *H* corresponds to the rarest subclone. DECODE thus finds the distribution of *θ*_*H*_ such that the SFS predicted by Eq. 1 best matches the empirical SFS from *inference A* and *inference B* subsamples (***step 3***, Fig. 1B).

To determine the parsimonious cluster configuration, DECODE tests the capacity of the inferred parameter distribution to predict the SFS from the *validation* subsample (***step 4***, Fig. 1B). The accuracy of the prediction, balanced against the model’s complexity as determined by cluster count *H*, is quantified with the Generalized Information Criterion (GIC) [Konishi and Kitagawa, 1996] (***step 5***, Fig. 1B). DECODE compares the GIC densities based on an increasing sequence of cluster counts. It selects the more complex result as the better model if its GIC is lower, and continues adding more clusters (***step 6***, Fig. 1B). Otherwise, it selects the model with the lower cluster count (***step 7***, Fig. 1B) as the most parsimonious decomposition (***step 8***, Fig. 1B). DECODE fits the data with and without a neutral tail, then selects the better model by GIC.

Two distinctions set DECODE apart from previous methodologies. First, the incorporation of mutation filtering strategies enables it to validate its results even for single-sample data. This reduces the risk for DECODE to either overfit or underfit the mutation clusters. Second, incorporating sample-specific sequencing coverage directly in the mathematical framework both enhances DECODE’s detection of overlapping clusters and ensures unbiased estimation for neutral tail power. Many existing methods assume mixture models for the SFS where each cluster density is characterized with parameters for the mean and dispersion [Carter et al., 2012, Roth et al., 2014, Miller et al., 2014, Deshwar et al., 2015, Caravagna et al., 2020]. Their criteria for selecting the parsimonious model thus strongly penalize decompositions with significantly overlapping clusters, to prevent overfitting due to the high parameter count. In contrast, the distribution for each cluster *h* in DECODE is defined only with its mean VAF *p*_*h*_, whereas its dispersion and overall shape are determined by the coverage distribution. We therefore expect that DECODE is more sensitive at detecting overlapping clusters, to be confirmed in the next sections. Furthermore, non-uniform coverage most significantly affects the low-VAF region in the SFS, which distorts the shape of the neutral tail [Dinh et al., 2020]. By accounting for this effect, DECODE is able to detect and characterize the tail more accurately and consistently.

### 2.2 DECODE outperforms existing methods in inferring tumor clonality and detecting neutral tail from the SFS

We first examined DECODE’s performance in inferring different aspects of tumor heterogeneity from single genomic samples by utilizing the ICGC-TCGA DREAM Somatic Mutation Calling Tumor Heterogeneity and Evolution Challenge. The DREAM Challenge consisted of 51 synthetic tumors, simulated under different assumptions of clonality, copy number aberrations, cellularity, sequencing depth and spatial growth that mirror real tumors [Salcedo et al., 2020]. The group compared 31 subclonal reconstruction algorithms and five reference methods based on the accuracy of the inferred sample purity (***task 1A***), subclone count (***task 1B***), cellular prevalences and mutation counts of each subclone (***task 1C***), deterministic and probabilistic assignments of mutations to subclones (***task 2A-B***), and deterministic and probabilistic subclonal phylogeny (***task 3A-B***) [Salcedo et al., 2025]. As DECODE was designed to cluster mutations with similar copy numbers, we limited the analysis to predominantly diploid samples (*n* = 29). DECODE currently does not support the prediction of subclonal phylogenies, so our comparison focused on ***tasks 1A-2B***. Since DREAM concentrated on the accuracy of mutation clustering, we implemented DECODE without the tail compartment, and compared its performance against the 36 software packages benchmarked in [Salcedo et al., 2025].

DECODE ranks among the top methods for purity inference (Fig. 2A). Its mean score across samples, normalized to the range between best and worst software akin to the DREAM protocol [Salcedo et al., 2025], is 7% worse than PhyloWGS, the leading method [Deshwar et al., 2015]. DECODE’s average score is most impacted by sample S5, where the synthetic tumor contains two independent clones, both at 40% cellular prevalence (i.e., VAF 20%), yielding an overall tumor purity of 80% (Supplementary Fig. 1A). However, DECODE’s predicted purity is based on the assumption that the truncal cluster in the SFS consists of mutations present in every tumor cell (Fig. 1A). Therefore, although it correctly identifies a truncal cluster at *p*_1_ = 0.2 in S5, DECODE incorrectly infers a purity of 2 · *p*_1_ = 40% for sample S5, similar to other approaches that do not take copy numbers into acount (Fig. 2A, Supplementary Fig. 1B, C). DECODE’s predicted purity is much closer to the true values for samples where cancer cells share a common ancestor, as we expect to be the case for many real tumors.

**Figure 2:**
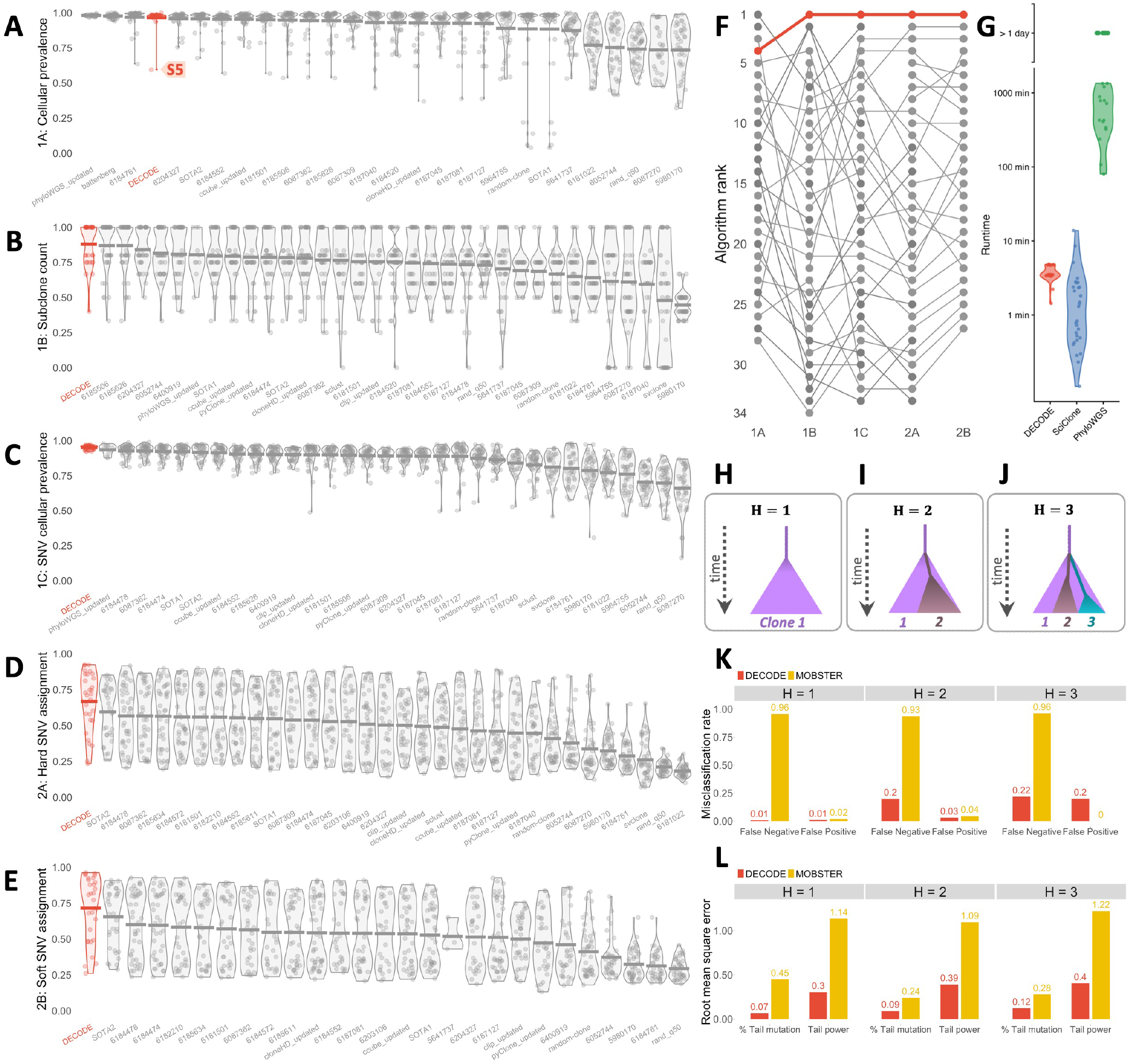
Synthetic benchmarking results. **A-G**: Results from benchmarking DECODE’s performance against 31 existing subclonal reconstruction methods analyzed in ICGC-TCGA DREAM Somatic Mutation Calling Tumor Heterogeneity and Evolution Challenge. **A-E**: DREAM score distributions for DECODE (red) and other algorithms (gray) in inferring cellular prevalence (**A**), subclone count (**B**), subclonal cellular prevalences and mutation counts (**C**), and deterministic and probabilistic mutation assignments (**D, E**). Each violin plot illustrates one algorithm’s scores across synthetic tumors (dots). Algorithms are ranked based on average scores across all samples (horizontal bars). **F**: Ranking of DECODE (red) and other algorithms (gray) across all DREAM tasks, based on average scores in each task. Each line connects available results from an algorithm. **G**: Comparison of computational runtime between DECODE, SciClone and PhyloWGS for analyzing DREAM samples. Each violin plot illustrates an algorithm’s runtime across synthetic tumors (dots). Runs that remained unfinished after 10^5^ seconds are marked as incomplete (“*>*1 day”). **H-L**: Results from benchmarking DECODE’s tail detection and characterization against MOBSTER. **H-J**: Schematics of clonal evolution models for simulations with *H* = 1 (**H**), 2 (**I**) and 3 subclones (**J**). **K, L**: Misclassification rates (**K**) and root mean square errors (RMSEs) of estimated tail power and mutation proportion (**L**) from DECODE and MOBSTER across the three simulation batches **H-J**.

DECODE’s inferences for subclone count and subclonal prevalences outperform algorithms benchmarked in the DREAM Challenge (Fig. 2B, C), with 2% and 5% improvements in normalized mean scores compared to previous best results, respectively. This seems unexpected, given that DECODE only analyzes mutations with the same ploidy, and therefore may overlook subclones associated with copy number variations (CNVs). The results suggest that DECODE is sensitive enough to detect CNV-driven subclonal expansions, given that they harbor sufficient mutations in their initiating cells to form an observable cluster in the SFS.

Compared with existing algorithms, DECODE performs particularly well in assigning mutations to individual subclones in both deterministic and probabilistic configurations (Fig. 2D, E), with 18% and 17% improvements compared to previous best methods, respectively. Therefore, we expect DECODE to accurately identify potential driver events for each subclone in single samples. Moreover, applications of DECODE in large-scale studies have the potential to reliably reveal frequent clonal and subclonal driver mutations.

Samples T3 and P5 in the DREAM Challenge provide some insight into DECODE’s performance (Supplementary Figs. 2, 3). The ground truth SFS for T3 consists of a truncal cluster at VAF *p*_1_ = 0.3 and a small cluster at *p*_2_ = 0.18, the latter obscured by a large cluster at *p*_3_ = 0.1 (Supplementary Fig. 2A). The substantial overlap among these clusters leads many existing methods to identify only the two dominant ones, which underestimates the heterogeneity in the sample (Supplementary Fig. 2B). In contrast, DECODE detects that the dispersions of these clusters, governed by the coverage distribution, are not wide enough to cover the intermediate VAF region. It thus correctly identifies three embedded clusters at frequencies *p*_1_ = 0.29, *p*_2_ = 0.19 and *p*_3_ = 0.1 (Supplementary Fig. 2C). On the other hand, sample P5 consists of a truncal cluster at *p*_1_ = 0.47, a subclonal cluster at *p*_2_ = 0.19 and a peak formed of false positives in the low-VAF region (Supplementary Fig. 3A). SciClone [Miller et al., 2014] infers five subclones, two of which overfit the truncal cluster (Supplementary Fig. 3B). DECODE accurately detects two clusters at *p*_1_ = 0.46 and *p*_2_ = 0.18, in addition to a third group at *p*_3_ = 0.05 consisting of the false positives (Supplementary Fig. 3C). Together, these two examples underscore DECODE’s enhanced accuracy in detecting overlapping clusters, reducing the risk of either underfitting or overfitting cancers’ heterogeneity.

**Figure 3:**
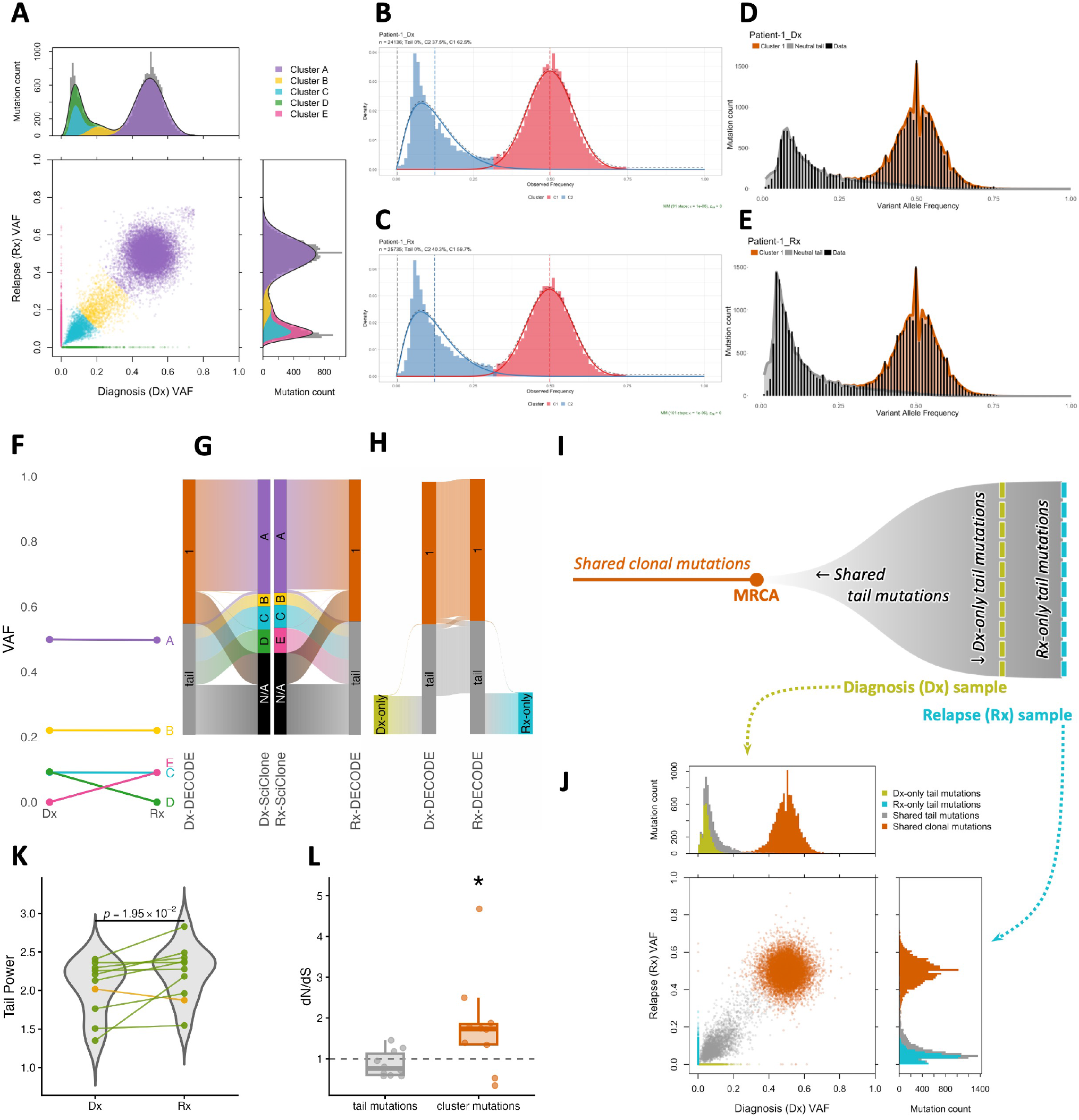
DECODE reveals neutral evolution in longitudinal acute myeloid leukemia. **A-J**: Mutation clustering for patient AML-1. **A**: SciClone’s joint clustering of mutations at diagnosis (Dx) and relapse (Rx). **B-E**: Deconvolution of mutations using MOBSTER (**B, C**) and DECODE (**D, E**) for each sample at diagnosis (**B, D**) and relapse (**C, E**). **F**: Mean VAFs between diagnosis and relapse of subclones inferred from SciClone. **G**: Alluvial diagrams of subclonal assignments for mutations at diagnosis (left) and relapse (right) between DECODE and SciClone. N/A: mutations without SciClone assignments. Widths are proportional to mutation fractions. **H**: Alluvial diagram of DECODE subclonal assignments for mutations at diagnosis and relapse. Outer bars contain mutations unique to each sample. **I**: Proposed model of evolution for patient AML-1. **J**: Synthetic multi-sample data simulated with TEMULATOR under the evolutionary model in **I. K**: Comparison of tail powers *α* at diagnosis and relapse inferred from DECODE. Lines connecting *α*’s for the same patients are highlighted in green if *α* is higher in Rx, and yellow otherwise. p-value from one-sided Wilcoxon test. **L**: dN/dS analysis of mutations assigned to tails and clusters in each patient (dots). *: p-value from one-sided Wilcoxon test < 0.05 (alternative: dN/dS *>* 1).

Notably, unlike most methods which excel on only a few objectives, DECODE performs strongly across all DREAM tasks (Fig. 2F). This is likely a result of its incorporation of sample-specific coverage in the SFS formulation and rigorous framework to select the parsimonious SFS deconvolution, which together enhance DECODE’s cluster detection. Across the 29 DREAM samples analyzed, SciClone [Miller et al., 2014], a popular clustering software, finishes in two minutes on average per sample (Fig. 2G). PhyloWGS [Deshwar et al., 2015], currently the state of the art (Fig. 2A-E), exceeds 1 day for 13 samples and requires 100 minutes or more in the remaining 16 samples (Fig. 2G). PhyloWGS’s long runtime is likely a result of its Markov chain Monte Carlo implementation, which is inherently difficult to parallelize, and its simultaneous analysis of both mutations and copy number aberrations, the latter not being utilized by DECODE or SciClone. DECODE is comparable to SciClone at roughly four minutes on average per sample, and is therefore applicable for large-scale genomic studies.

We further benchmarked DECODE’s detection and characterization of the SFS neutral tail against MOBSTER, currently the only other method that incorporates this component to our knowledge [Caravagna et al., 2020]. We simulated data comprising 1 − 3 clones (Fig. 2H-J). For each clone count, we created 1,800 synthetic tumors with tails (“T-type”) and 200 simulated samples without tails (“NT-type”). Each sample was simulated with purity, mutation burden, and sequencing coverage drawn at random from The Cancer Genome Atlas (TCGA) [Weinstein et al., 2013]. DECODE detects the neutral tail more reliably than MOBSTER, with lower false negative rates (i.e., missed tails in T-type samples) and false positive rates (i.e., spurious tails in NT-type samples) across all clone counts, with the exception of the false positive rate in *H* = 3 (Fig. 2K). This likely reflects MOBSTER’s tendency to detect the tail confidently only at high coverage (> 100×), and to be more conservative at the lower coverage typical of TCGA [Caravagna et al., 2020, Salcedo et al., 2025]. Furthermore, when restricted to each method’s true positives (i.e., T-type samples in which the tail is successfully detected), DECODE’s estimates of the tail power and the fraction of mutations belonging to the tail are also significantly more accurate than MOBSTER’s (Fig. 2L). This highlights the need to account for sample-specific coverage and mutation-calling biases, which substantially distort the SFS tail.

We next investigated which attributes of genomic data most strongly affect the accuracy of DECODE’s inferred tail power and mutation proportion (Supplementary Figs. 4-6). As can be expected, DECODE’s results tend to be more accurate for samples with higher purity, coverage depth and mutation count. Nevertheless, only two properties consistently affect DECODE’s tail characterization at a significant level across all clone counts. The first is cluster count error: over- or under-estimating the clonality distorts the tail compartment, leading to inaccurate tail parametrization (Supplementary Figs. 4-6). However, this is unlikely to affect many clinical applications, as DREAM results indicate that DECODE’s inferred clonality for realistic samples is generally reliable (Fig. 2B). The second significant factor is entropy in the tail region (Supplementary Figs. 4-6). High entropy, caused by low-VAF subclones obscuring the tail, renders tail parameters non-identifiable. Hence, for samples with low purity, high clonality or rare subclones, we expect DECODE’s tail characterization to be less reliable.

Together, the results from the DREAM Challenge and our SFS tail benchmark study suggest that DECODE is well positioned to become a new standard in characterizing tumor clonality and evolution mode from individual genomic samples, with potential for clinical applications.

### 2.3 Incorporating the neutral tail enhances the reconstruction of clonal evolution in acute myeloid leukemia

While DECODE was designed to decompose single genomic samples, we also explored its utility for multi-sample data by analyzing paired acute myeloid leukemia (AML) whole-genome sequences at diagnosis (Dx) and relapse (Rx) from 10 patients in [Shlush et al., 2017]. Multi-sample deconvolution with SciClone [Miller et al., 2014] infers 4-7 subclones per patient (Fig. 3A, Supplementary Figs. 7-15). Analyzing each of the 20 samples individually, MOBSTER distinguishes 2-3 subclones per sample (Fig. 3B-C, Supplementary Figs. 7-15) but detects the neutral tail in only one sample, which was sequenced at 137× (Supplementary Fig. 14B). MOBSTER’s inability to identify the tail in 19*/*20 cases is likely because the sequencing depths of these samples (median 54×, range 45 − 70×) are significantly lower than the *>* 90 − 100× threshold recommended by its authors. Meanwhile, DECODE identifies neutral tails in all 20 samples, in addition to 1 − 3 subclones per sample (Fig. 3D-E, Supplementary Figs. 7-15). We investigated the evolution models extrapolated from DECODE’s tail-aware fits, in comparison to SciClone’s joint clustering results.

In Patient 1, SciClone detects one subclone present only at diagnosis, another subclone specific to relapse, and three subclones shared at both time points (Fig. 3A). Intriguingly, the mean VAFs of the shared subclones remain unchanged from diagnosis to relapse (Fig. 3F). The implication of clonal selection equilibrium may be biologically questionable, given that the intervening treatment resulted in clinical and morphological remission in this patient [Shlush et al., 2017]. In contrast, DECODE determines that the mutations with VAF ≈ 0.5 in both samples (i.e., SciClone’s cluster A) form the truncal cluster, while the other two shared subclones and both Dx- and Rx-specific subclones inferred from SciClone (i.e., clusters B, C, D and E) instead form the neutral tails in each sample (Fig. 3G).

DECODE’s truncal cluster mutation assignments are highly consistent across samples, despite each sample being decomposed independently (Fig. 3H). Likewise, the majority of shared mutations with lower VAFs are assigned to the tail in both samples, with the mutations specific to each time point making up the remainder of each tail. We therefore hypothesize that the diagnosis sample consists of cells proliferating at similar rates, and the relapse sample simply reflects a continuation of their expansion following a treatment-induced bottleneck (Fig. 3I). Synthetic multi-sample data, simulated under this evolution model with TEMULATOR [Heide, 2022] (Fig. 3J) are indeed consistent with the joint SFS pattern observed in Patient 1’s data (Fig. 3A). In this evolution model, the mutations present in the primary tumor’s MRCA form a shared clonal cluster, centered at VAFs determined by the cellular prevalence at each time point. Mutations arising soon after the MRCA exist in relatively high cell fractions, thus they comprise a portion of the diagnosis SFS tail that persists at relapse. The shared tail mutations therefore form a cloud in the joint SFS, distributed along the diagonal from the low-VAF corner to the shared clonal cluster. In contrast, late-arising mutations exist in low cell fractions at diagnosis and are prone to imminent extinction, contributing a Dx-specific tail at lower VAFs than the shared tail. Finally, mutations emerging after diagnosis constitute the Rx-specific SFS neutral tail. This evolution model is consistent with the original publication, which found that the diagnosis and relapse blasts in this patient are highly related [Shlush et al., 2017]. The joint SFS of 6*/*10 patients mirror Patient 1’s; in each case, DECODE’s deconvolutions consist only of neutral tails and truncal clusters (Supplementary Figs. 8, 11, 12, 13, 15). We therefore hypothesize that the evolution of these tumors follows the model in Fig. 3I. DECODE detects subclones in the remaining 4 cases, three of which exhibit higher clonality at relapse (Supplementary Figs. 7, 9, 10).

Previous work in population genetics has revealed that the SFS tail power *α* is an indicator of ongoing tumor expansion, with *α* = 1 in cell populations of constant size [Watterson, 1975] and *α* = 2 in exponentially expanding populations [Griffiths and Tavaré, 1998]. DECODE results demonstrate an increase in tail power at relapse in 9*/*10 AML patients (Fig. 3K), suggesting that cancer cells expand more aggressively at relapse than at diagnosis, even in patients with no detectable subclones. Finally, dN/dS analysis confirms that mutations that DECODE assigns to the SFS tail are consistent with neutral evolution (i.e., dN/dS ≈ 1) while the variants assigned to mutation clusters indicate positive selection (i.e., dN/dS *>* 1), further corroborating DECODE’s detection of neutral tails and clonality in the SFS (Fig. 3L).

This study underscores three advantages of applying DECODE to analyze cancer genomic data. First, it is able to identify neutral tails in the SFS even for samples with low sequencing depths, thanks to our correction for coverage distribution and mutation filtering. Second, by accounting for mutations in the SFS tail, it avoids inferring spurious subclones from neutrally accumulating passenger mutations. This often results in simpler and more biologically plausible reconstructions of tumor evolution. Finally, DECODE’s inferred tail powers provide an orthogonal measure for characterizing single cancer samples, independent from clonality. SFS tail power reflects the ongoing tumor expansion mode, which has been observed to be a strong biomarker for patient survival in time-series experiments [Stein et al., 2008]. By enabling the estimation of tumor expansion rates from single genomic samples, DECODE has the potential to support clinical decision-making in settings where serial or multi-region sampling is impractical.

### 2.4 DECODE infers clonality, tail power and sample purity accurately and consistently for pan-cancer samples

We implemented DECODE to analyze pan-cancer whole-genome sequencing (WGS) samples from The Cancer Genome Atlas (TCGA) [Weinstein et al., 2013]. We first examined the consistency of DECODE’s results across same-patient biological replicates, sampled from different tumor regions across multiple cancer types. DECODE’s assignment of mutations to the tail or specific SFS clusters is more concordant across 57 multi-region WGS pairs, compared to SciClone’s subclonal assignments (Fig. 4A; median adjusted Rand index (ARI): 0.19 in DECODE, 0.09 in SciClone). Because samples from the same patient are expected to share a broadly common evolutionary history, the concordance indicates that DECODE recovers reproducible clonal structure rather than sample-specific noise.

**Figure 4:**
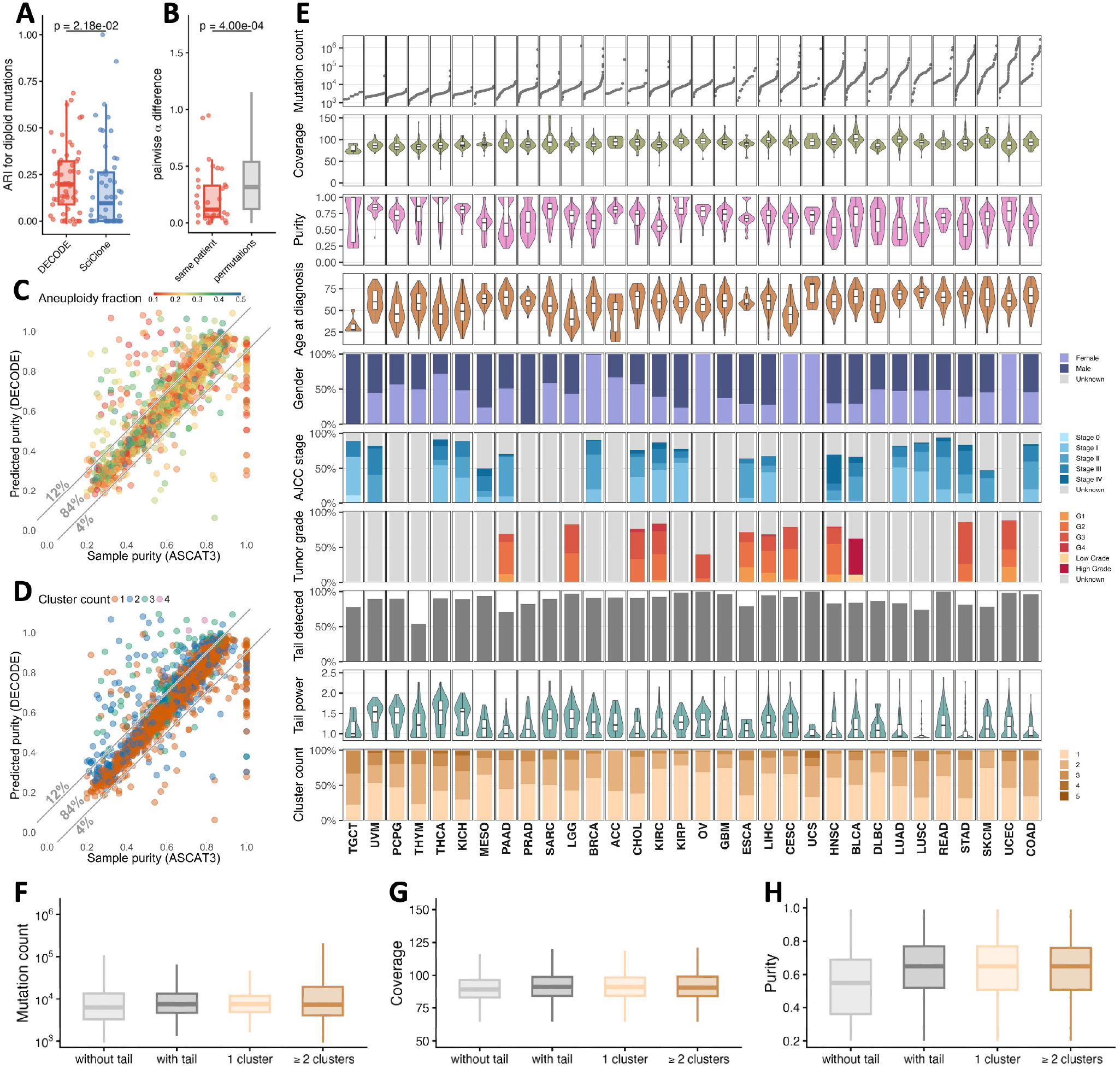
Deconvolutions of pan-cancer samples in TCGA. **A**: Comparison of ARI scores for deconvolutions of same-patient biological replicates in TCGA between DECODE and SciClone. p-value from one-sided Wilcoxon test. **B**: Comparison of differences in DECODE-inferred tail powers among same-patient biological replicates and permutated samples. p-value from permutation test. **C, D**: Comparison of sample purities estimated by ASCAT3 against those predicted from DECODE-inferred truncal clusters, with colors indicating sample aneuploidy fraction (**C**) or DECODE-inferred cluster count (**D**). ASCAT3 assigns purity = 1 for samples where aneuploidy fractions are too low for accurate estimation. Diagonal lines demarcate correctly predicted purities (error < 0.1) from over- and under-estimates; percentages indicate the proportion of samples in each group. **E**: Summary of tumor characteristics and DECODE results across TCGA cancer types, ordered by median mutation burden. Top to bottom: sample-specific mutation burdens; whole-genome sequencing depths; ASCAT3-inferred sample purity; patient ages at diagnosis; patient genders; tumor stages per the American Joint Committee on Cancer (AJCC) system; tumor grades; proportions of samples in which DECODE detected neutral tails; DECODE-inferred tail powers; and DECODE-inferred cluster counts. **F-H**: Distributions of mutation burden (**F**), sequencing depth (**G**) and sample purity (**H**) across all samples, stratified by whether DECODE detected a tail and its inferred cluster count.

Additionally, among 35 pairs of biological replicates where DECODE detects a tail in both samples, the inferred tail powers agree significantly more closely within the same patients than across permuted pairs (Fig. 4B). Given that the AML study establishes the importance of the tail power as an indicator of tumor severity (Fig. 3K), this agreement further reaffirms the robustness of the tail power as a measure of tumor expansion.

As in the DREAM Challenge (Fig. 2A), cellular prevalence can be predicted from the truncal cluster VAF. In 84% of the TCGA samples with aneuploidy fraction *>* 0.1 (*n* = 1,966/2,333), the predictions are within 10% of the sample purities inferred from ASCAT3 [Van Loo et al., 2010] (Fig. 4C, D). Among samples where DECODE’s prediction error exceeds 10%, roughly one in four are underestimates. This mainly results from the low level of aneuploidy in these samples, due to which ASCAT3 and other copy number-based methods cannot estimate purity accurately (Fig. 4C). DECODE overestimates the cellular prevalence in 12% of the samples, indicating that alternative fits with fewer clusters and smaller truncal cluster VAFs would have been more appropriate for these samples (Fig. 4D). We note that errors in allele-specific copy number calling can introduce spurious high-VAF clusters in the SFS. One such case is copy-neutral loss of heterozygosity miscalled as heterozygous: in a sample of purity *ρ*, truncal mutations on the retained allele then appear at VAF *ρ* rather than the expected *ρ/*2, mimicking a subclone at high cellular prevalence and rendering the maximum cluster VAF an unreliable predictor for purity. Overall, copy number-based methods remain the most robust approach for estimating sample purity from genomic data. However, for samples with low aneuploidy where these methods have limited signal (e.g., 45% samples in TCGA have aneuploidy fraction ≤ 0.1), the strong agreement with ASCAT3 supports DECODE as a complementary route to estimating cellular prevalence.

We evaluated DECODE’s deconvolutions of primary tumors in TCGA (*n* = 4,058 patients across 32 cancer types). DECODE detects the neutral tail in 89% of the samples, with cancer-specific proportions ranging from 54% in thymomas (THYM) to 100% of ovarian cancers (OV), rectum adenocarcinomas (READ) and uterine carcinosarcomas (UCS) (Fig. 4E). In addition, it infers the existence of multiple mutation clusters in 47% of the patients (*n* = 1,898/4,058), with the proportion of samples with two or more clusters ranging from 22% in kidney renal papillary cell carcinomas (KIRP) to 78% in testicular germ cell tumors (TGCT). The inferred tail powers fall between 1 and 2 for the majority of patients (67%, *n* = 2,433/3,625), with only 1% of samples (*n* = 51/3,625) having tail powers higher than 2. As noted above, this indicates that most tumors occupy a continuum between stable size (*α* ≈ 1) and exponential expansion (*α* ≈ 2).

DECODE’s inference of clonality does not seem to be affected by mutation burden, sequencing coverage or sample purity (Fig. 4F-H). Its detection of neutral tail in the SFS likewise does not depend on variant count or read depth (Fig. 4F-G). However, samples where DECODE cannot identify a tail have significantly lower purity than those where it does (Fig. 4H). This is because, at low cellular prevalence, the mutation clusters are compressed into the low-VAF region in the SFS where the tail resides. The lack of a clear boundary between tail and clusters then leads DECODE to prefer a more parsimonious fit without a tail.

### 2.5 DECODE clonality stratifies overall survival in low-grade glioma

We assessed whether DECODE-inferred clonality is associated with patient outcome. We defined TCGA cohorts by cancer type, and further stratified each cancer type by histological stage or tumor grade. Stratifying patients in each cohort into two subgroups by the median DECODE-inferred cluster count reveals significant survival differences in two cohorts (Fig. 5A). The first is the primary low-grade glioma (LGG) cohort, where patients with polyclonal tumors (two or more subclones) survive significantly longer than those with monoclonal tumors (Fig. 5B; median survival time: 4.1 years in samples with one cluster, 10.9 years in samples with two or more clusters). Similarly, LGG grade 3 patients with multiple detected subclones survive on average roughly three times longer compared to those with clonally homogeneous tumors (Fig. 5C; median survival time: 2.6 years in samples with one cluster, 7.8 years in samples with two or more clusters). The fact that cluster count is clinically meaningful within the LGG grade 3 cohort indicates that for some cancer types, DECODE-inferred clonality may provide prognostic information beyond what histology and staging capture.

**Figure 5:**
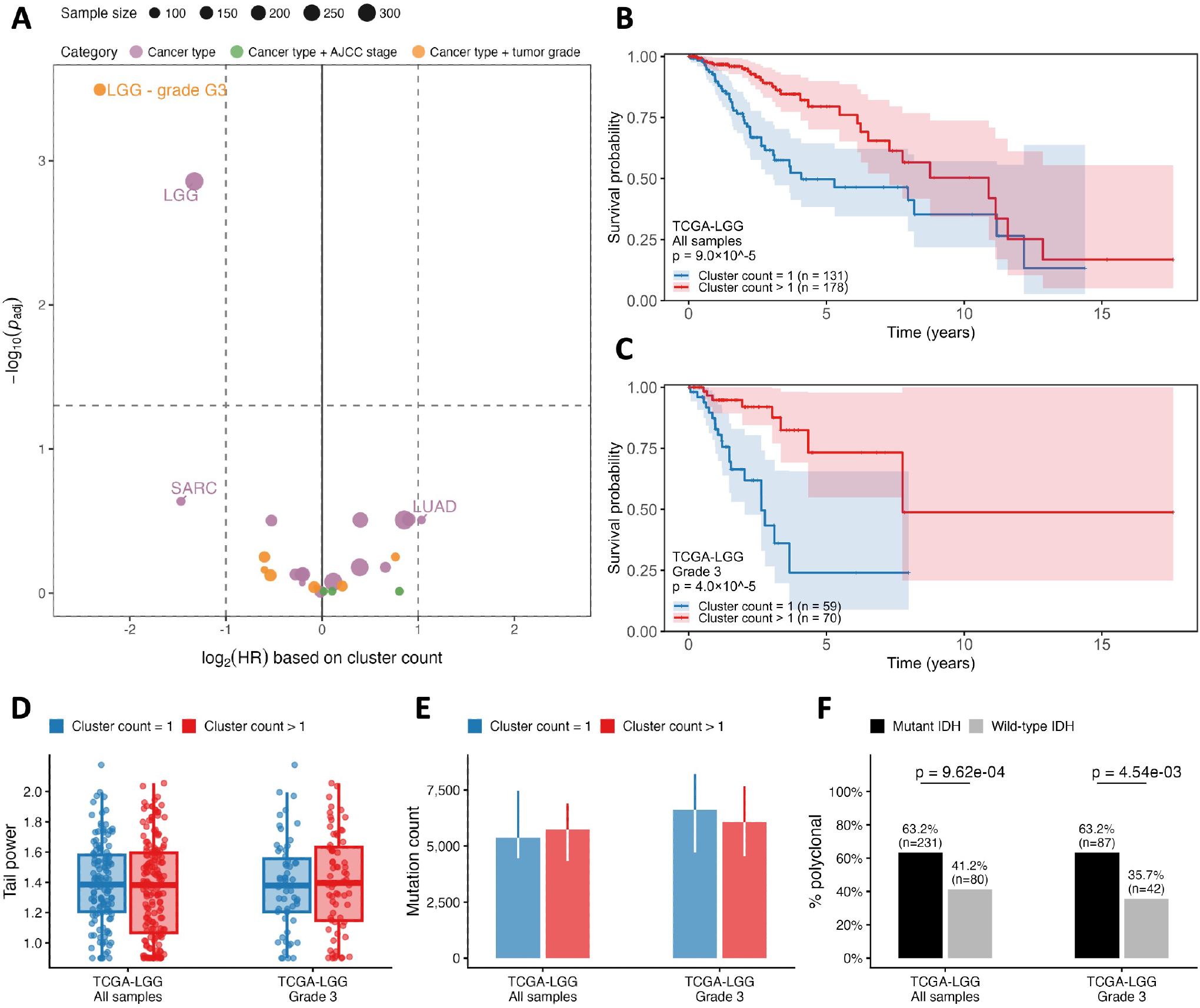
Prognostic value of DECODE-inferred clonality in TCGA. **A**: Volcano plot summarizing the association between cluster count and overall survival across TCGA cohorts, defined by cancer type alone, cancer type with AJCC pathological stage, or cancer type with tumor grade. Samples within each cohort were split at the median cluster count; the x-axis shows the hazard ratio (HR) between the resulting subgroups, and the y-axis shows the Benjamini–Hochberg adjusted log-rank p-value *p*_adj_ comparing their Kaplan–Meier curves. Dashed lines mark *p*_adj_ = 0.05, and HR = 2 and 1*/*2. **B, C**: Kaplan–Meier overall survival curves for the TCGA lower-grade glioma (LGG) cohort, comprising all samples (**B**) or only grade 3 tumors (**C**), partitioned at the median cluster count (= 1 in both). p-values from log-rank tests comparing subgroups in each cohort. **D, E**: DECODE-inferred tail powers (**D**) and mutation burdens (**E**) in the LGG and LGG grade 3 cohorts, partitioned at the median cluster count (= 1). **F**: Proportions of samples with cluster count *>* 1 in the LGG and LGG grade 3 cohorts, partitioned by IDH status. p-values from two-sided Fisher’s exact test.

We investigated the biological factors driving the survival differences uncovered by DECODE clonality in LGG. There is no significant difference in tail power or mutation count between patient subgroups in either the LGG or LGG grade 3 cohort (Fig. 5D-E), suggesting that the survival disparity is unlikely to reflect differences in tumor expansion mode or mutation burden. However, DECODE detected multiple subclones significantly more often in IDH-mutant than in IDH-wildtype tumors in both cohorts (Fig. 5F). IDH-mutant LGGs are known to grow more slowly than IDH-wildtype tumors [van den Bent et al., 2024] and to confer substantially longer overall survival [The Cancer Genome Atlas Research Network, 2015]. We therefore hypothesize that IDH-mutant LGG’s longer evolutionary timescale gives subclones more time to diversify and reach detectable VAF, potentially under near-neutral dynamics [Williams et al., 2016], which would account for the strong association between higher ITH and favorable outcome (Fig. 5B-C). Although the association between IDH mutation and elevated ITH has not been reported in LGG to our knowledge, mutant IDH has been linked to increased clonality in intrahepatic cholangiocarcinoma [Xiang et al., 2021] and acute myeloid leukemia [Sirenko et al., 2025].

## 3 Discussion

DECODE is a novel mutation clustering method that builds on our previous work in population genetics and parameter inference [Dinh et al., 2020, 2025]. The synthetic tests showed that DECODE can detect and characterize the SFS neutral tail accurately. Furthermore, DECODE estimates multiple aspects of tumor clonality more reliably and consistently than current approaches (Fig. 2). Incorporating the tail in the SFS simplifies the decomposition of many acute myeloid leukemias, yielding a more biologically realistic model of how these cancers evolve toward relapse (Fig. 3). Applied to TCGA pan-cancer data, DECODE reveals an SFS tail in the majority of samples across tumor types (Fig. 4), and its clonality estimates reveal a correlation between ITH and survival in low-grade glioma (Fig. 5). Overall, DECODE’s clonality and tail-power inferences provide two complementary axes for estimating tumor clonality and expansion mode from genomic data, both of which have been shown to be important biomarkers for patient outcome [Swanson et al., 2008, Ferté et al., 2014, Andor et al., 2016, Caswell and Swanton, 2017, Dagogo-Jack and Shaw, 2018].

Despite these advantages, DECODE has several areas for improvement. First, because it relies on the VAF-based SFS, DECODE’s clonality inference is valid only for variants sharing the same copy-number state. It is therefore suitable to analyze most diploid tumors, but not those with increased chromosomal instability. Second, DECODE was designed to analyze individual samples. Unlike PhyloWGS and SciClone [Deshwar et al., 2015, Miller et al., 2014], among others, it currently cannot jointly analyze multi-region or longitudinal samples from the same patient. However, our AML study showed that DECODE’s single-sample results can be merged to reconstruct the overall clonal evolution across multiple samples (Fig. 3). Finally, DECODE does not explicitly infer the phylogenetic relationships between subclones. Applying constraints such as the pigeonhole principle, integrating tree-building approaches [Deshwar et al., 2015], or extending DECODE to multi-sample data may enable reconstruction of the subclonal phylogeny.

## Methods

### A DECODE’s framework

#### A.1 Expected site frequency spectrum given coverage distribution and mutation filtering

We assume that a tumor sample consists of *H* subclones, and that the fraction *c*_*h*_ of sampled cells belonging to subclone *h* share the *ω*_*h*_ mutations present before the subclone’s most recent common ancestor (MRCA) (Fig. 1A). We denote the ancestral clone with *h* = 1 from which other subclones *h* = 2, … , *H* originate, so that *c*_1_ constitutes the cellular prevalence (i.e., sample purity). Assuming diploidy, the expected number of mutations present in *k* copies among *n* alleles from a sample of *n/*2 cells, following results in population genetics [Griffiths and Tavaré, 1998, Durrett, 2013, Dinh et al., 2020], is:

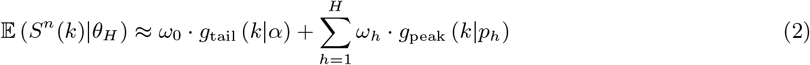

where *θ*_*H*_ = {*α, p*_1_, … , *p*_*H*_ , *ω*_0_, … , *ω*_*H*_} are the parameters to be inferred. Mutations in cluster *h* follow a binomial distribution with mean variant allele frequency (VAF) *p*_*h*_ = *c*_*h*_*/*2 [Tarabichi et al., 2021]:

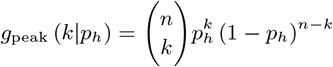

The *ω*_0_ mutations in the “foreground” occur neutrally in all subclones, following a tail distribution with power *α*:

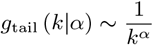

for *k* = 1, … , *n*· *p*_1_. Previous works have shown that for well-mixed samples, the neutral mutation tail is of power *α* = 1 in a population of constant size [Watterson, 1975], and *α* = 2 in an exponentially expanding population [Griffiths and Tavaré, 1998]. Different modes of tumor growth, spatial effects and other confounding factors can have an impact on *α*.

We previously developed the formula for the expected observed SFS that accounts for the sequencing coverage and data filtering [Dinh et al., 2020], relevant details of which are described here. We define the coverage as the probability mass function *φ*, such that *φ*(*r*) is the probability that a site is covered by a total of *r* reads. Because DNA-seq samples typically contain ≥ 10^5^ cells and are sequenced with mean coverage < 100×, we assume that the *r* reads originate from different cells. For a mutation present in *k* alleles in the sample, its exact VAF is *k/n*, therefore the number of variant reads follows *z*|*r* ∼ Binomial(*r, k/n*). The expected mutation count with observed frequency *f* ∈ (*f*_1_, *f*_2_], for 0 ≤ *f*_1_ *< f*_2_ ≤ 1, is then:

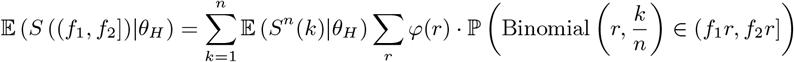

where ℙ 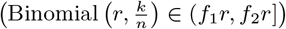 is the probability that a mutation covered by *r* reads with true VAF *k/n* has observed VAF *f* = *z/r* ∈ (*f*_1_, *f*_2_].

The sequencing data can be represented as *D* = {(*z*_*i*_, *r*_*i*_)}_*i*=1,2,…_, where the *i*th mutation is observed in *z*_*i*_ variant reads among *r*_*i*_ total reads. With thresholds *L, M* ≥ 0 for variant and total read counts, respectively, the filtered data is *D*^[*L,M*]^ = {(*z*_*i*_, *r*_*i*_) ∈ *D* : *z*_*i*_ ≥ *L, r*_*i*_ ≥ *M*}. The observed SFS 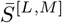 can then be tabulated from {*z*_*i*_*/r*_*i*_ : (*z*_*i*_, *r*_*i*_) ∈ *D*^[*L,M*]^} with bin size *b*. The expected observed SFS, accounting for the mutation filtering, is:

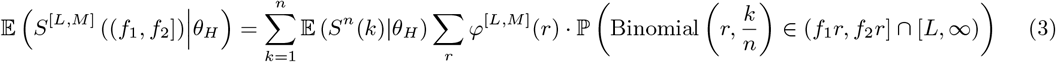

for each bin (*f*_1_, *f*_2_] = (0, *b*] , (*b*, 2*b*] , … The coverage distribution can be approximated directly from normalized counts of mutations with given total read counts in the filtered mutation data:

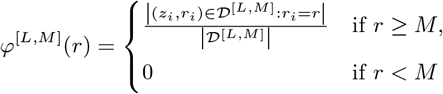

where | ∗ | counts the mutations in a dataset. Together, Eqs. (2) and (3) form a connection between the exact but unobserved SFS, which contains signals for the neutral tail and mutation clusters, and the expected observed SFS, which accounts for sample-specific sequencing coverage and mutation filtering. By default, DECODE assumes a sample with *n* = 1000 alleles and the SFS is computed with *N*_bins_ = 100 bins of size *b* = 0.01.

#### A.2 Inferring parameters for the neutral tail and mutation clusters

Assuming that the cluster count *H* is known, DECODE seeks *θ*_*H*_ = {*α, p*_1_, … , *p*_*H*_ , *ω*_0_, … , *ω*_*H*_} that best fits the observed mutational data. To ensure reliability, it analyzes data with two distinct filtering strategies (*L*^[*A*]^, *M*^[*A*]^) and (*L*^[*B*]^, *M* ^[*B*]^) simultaneously. By default, DECODE selects 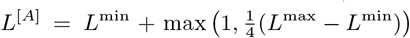 and 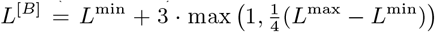, where *L*^min^ and *L*^max^ are the minimum and 95^*th*^ percentile of the variant read counts observed in the data, respectively. These thresholds filter out the mutations with the fewest variant reads, as they are most affected by mutation calling biases. Furthermore, increasing *L* changes the shape of the SFS, especially the neutral tail which mainly exists in the low-VAF region. The total read count thresholds *M* ^[*A*]^ and *M* ^[*B*]^ are then chosen such that the correspondingly filtered data subsets *D*^[*A*]^ and *D*^[*A*]^ contain roughly the same numbers of mutations, to avoid overfitting either subsample. DECODE is therefore required to fit the distinct SFS resulting from both (*L*^[*A*]^, *M* ^[*A*]^) and (*L*^[*B*]^, *M* ^[*B*]^) well, hence increasing the robustness of its results.

Following Eqs. (2)-(3), the expected number of tail mutations observed in *D*^[*A*]^ is 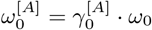, where

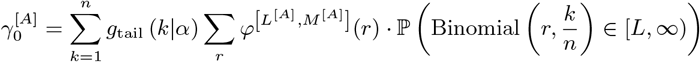

is a constant. Similarly, the expected number of mutations in cluster *h* in *D*^[*A*]^ is 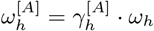, where

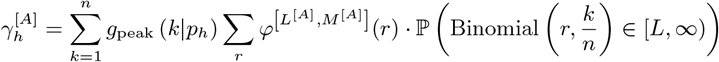

Therefore, inferring *θ*_*H*_ is equivalent to estimating 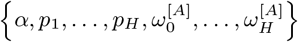, which is more practical because 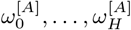 are naturally limited by the filtered mutation count |*D*^[*A*]^| , providing unbiased bounds for parameter estimation. For a given parameter set, the true mutation counts can be back-calculated as 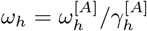.

To estimate 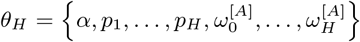, we implement Approximate Bayesian Computation sequential Monte Carlo via distributional random forests (ABC-SMC-DRF), a parameter inference method that we recently introduced which integrates distributional random forests (DRFs) in the framework of Approximate Bayesian Computation sequential Monte Carlo (ABC-SMC) [Dinh et al., 2025]. On the one hand, by implementing DRF, ABC-SMC-DRF circumvents the selection of statistical distance functions and tolerance thresholds in traditional ABC methods, which may affect the result’s accuracy. On the other hand, by being embedded in the ABC-SMC framework, it focuses on the most likely regions in the parameter space iteratively, thus converging to the posterior distribution faster with less computation.

DECODE implements a modified version of ABC-SMC-DRF to accommodate cluster reordering. In the first iteration, 1000 samples of *θ*_*H*_ are drawn from the prior distribution:

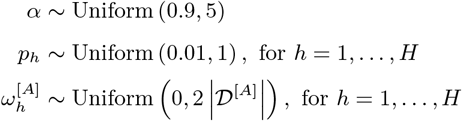

For each *θ*_*H*_ , the true mutation counts *ω*_0_, … , *ω*_*H*_ are computed, then the observed SFS is predicted from Eq. (3) for thresholds (*L*^[*A*]^, *M* ^[*A*]^) and (*L*^[*B*]^, *M* ^[*B*]^). The relative error for each SFS prediction ∗ = *A, B* compared to the correspondingly filtered SFS is defined as

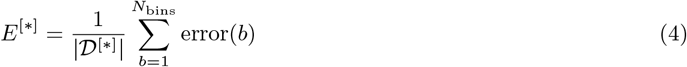

where

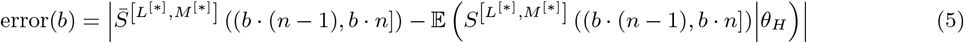

is the error in bin *b*. The total error for parameter set *θ*_*H*_ is then *E* = *E*^[*A*]^ + *E*^[*B*]^. DECODE trains a DRF on the 1000 particles of *θ*_*H*_ and corresponding errors, then derives weights for the particles to minimize the error [Cévid et al., 2022]. Finally, it reorders the pairs 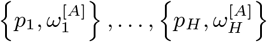 in each *θ*_*H*_ such that *p*_1_ *>* · · · *> p*_*H*_.

In the second iteration, DECODE samples 1000 new *θ*_*H*_ ‘s from the weighted particles in iteration 1. It then perturbs each parameter set with Gaussian kernels, where each kernel’s variance is twice the corresponding parameter’s empirical variance in iteration 1. This adaptive perturbation scheme has been shown to be unbiased and efficient in the context of ABC-SMC [Beaumont et al., 2009]. The new particles’ weights are then estimated with DRF and the clusters in each particle are sorted, similarly to iteration 1. DECODE continues until iteration 10, and the final weighted particle set approximates the posterior distribution for 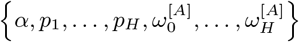.

To ensure stability and accuracy, DECODE also performs a guided parameter inference if the results for *θ*_*H*−1_ are available. It guesses where cluster *H* resides by finding the bin *b* with central VAF 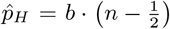 where the average error from Eq. (5) for *H* − 1 clusters is maximal, and posits that the mutation count for cluster *H* in *D*^[*A*]^ is approximately 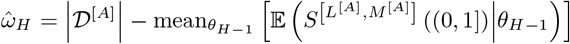, the average residual in the data compared to the SFS predicted with the previously inferred *θ*_*H*−1_ particles. The prior distribution for this inference is then

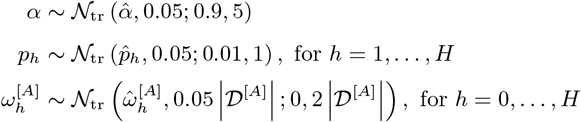

where 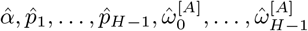 are the average estimates from *θ*_*H*−1_, and *N*_tr_(*µ, σ*; *a, b*) is the normal distribution with mean *µ* and variance *σ*^2^, truncated to within (*a, b*). DECODE then proceeds to estimate *θ*_*H*_ with the modified ABC-SMC-DRF procedure, similar to the inference from uniform prior distributions. Finally, the posterior distributions from uninformed and informed inferences are compared, and the result with lower average error is selected as DECODE’s final estimate for *θ*_*H*_.

#### A.3 Parsimonious model selection

DECODE seeks the smallest cluster count *H* such that *θ*_*H*_ captures the observed mutational data. To ensure robustness, DECODE examines the SFS predicted by the inferred *θ*_*H*_ for a third filtering strategy (*L*^[*V*]^, *M* ^[*V*]^) , where 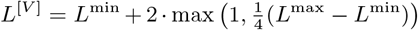 and *M* ^[*V*]^is the minimum total read count observed in the data. As *L*^[*A*]^*< L*^[*V*]^*< L*^[*B*]^, DECODE can validate whether the *θ*_*H*_ distribution inferred for filters (*L*^[*A*]^, *M*^[*A*]^) and (*L*^[*B*]^, *M* ^[*B*]^) can predict the novel SFS shape extracted from (*L*^[*V*]^, *M* ^[*V*]^). For each *θ*_*H*_ particle, we define the Generalized Information Criterion [Konishi and Kitagawa, 1996]:

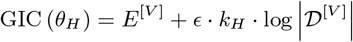

where *E*^[*V*]^is the relative error of the SFS prediction from Eq. (4), and *ϵ* = 0.0015 calibrates the penalty of parameter count *k*_*H*_ = 2*H* + 2*ϵ*_tail_, relative to the mutation count |*D*^[*V*]^| in the validation SFS subset, and *ϵ*_tail_ ∈ [0, 1] is the user-defined weight for neutral tail parameters.

After successive inferences with *H* − 1 and *H* clusters, DECODE quantifies whether the more complex model is justified by the improvement in prediction quality with a one-sided Wilcoxon rank sum test, where the null hypothesis is that median {GIC(*θ*_*H*−1_)} ≤ median {GIC(*θ*_*H*_)}. DECODE selects the model with *H* clusters and continues to infer *θ*_*H*+1_ if the p-value < 0.01, otherwise it determines that the model with *H* − 1 clusters is the best fit for the data.

If there are rare subclones in the sample, the low-VAF clusters may obscure the tail, leading to unreliable SFS tail characteristics. Therefore, similar to MOBSTER [Caravagna et al., 2020], DECODE also fits the data with only mutation clusters, where the parameter set consists of 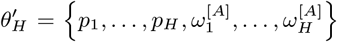. The inference of 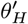 distributions and selection of the parsimonious model are similar to DECODE routines with tail. DECODE chooses the overall best model between the selected results with *H* clusters and a tail, and *H*′ clusters with no tail, with a one-sided Wilcoxon rank sum test for whether 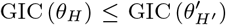. The model with *H*′ clusters and no tail is selected if the p-value < 0.01, otherwise DECODE chooses the model with *H* clusters and a tail. In practice, we find *ϵ*_tail_ = 0.3 to provide a good balance between ascertaining the tail parameters and selecting the clusters-only fit when the tail is too obscured.

### B Synthetic testing

#### B.1 ICGC-TCGA DREAM Challenge

We tested DECODE’s performance using the synthetic benchmark from the ICGC–TCGA (International Cancer Genome Consortium – The Cancer Genome Atlas) DREAM Somatic Mutation Calling Tumor Heterogeneity and Evolution Challenge [Salcedo et al., 2020]. The dataset consists of 51 synthetic tumors, created to mimic real cancer samples under various assumptions of clonality, copy number aberrations, cellularity and sequencing depth, while also accounting for the tumors’ spatial growth. The authors simulated tumor BAM files with BAMSurgeon [Ewing et al., 2015], then identified somatic single-nucleotide variants (SNVs) and copy number aberrations (CNAs) with Genome Analysis Toolkit (GATK) MuTect [Cibulskis et al., 2013] and Battenberg [Nik-Zainal et al., 2012], respectively.

As DECODE is designed to analyze mutations with the same copy number background, we limited the samples to be analyzed by DECODE to those where (a) there are ≥ 50 diploid mutations, and (b) the diploid mutations constitute ≥ 60% of all mutations. Of the 30 retained samples, one (sample P2) contains 123,969 diploid mutations. Although DECODE successfully deconvoluted this sample, the prohibitive memory demand for the co-clustering matrix (≈ 270 GB) prevented us from scoring ***task 2B***. Therefore, P2 was excluded from our analysis, and we reported the results for the remaining 29 synthetic tumors. Following DREAM protocol, subchallenge scores were reported as NA for the samples that we excluded.

For each sample, the input data for DECODE consisted of all diploid and heterozygous mutations (major CN = minor CN = 1). Because the DREAM Challenge does not account for the SFS tail, we implemented DECODE without a tail. The result from DECODE thus consisted of cluster count *H*, the distribution of *θ*_*H*_ = *p*_1_, … , *p*_*H*_ , *ω*_1_, … , *ω*_*H*_ , and the expected SFS {E (*S*^[*]^(*b*)|*θ*_*h|H*_), bin *b* = 1, … , *N*_bins_} for each cluster *h* = 1, … , *H* with parameters *θ*_*h*|*H*_ = {*p*_*h*_, *ω*_*h*_} under filters ∗ = (*L*^[*A*]^, *M* ^[*A*]^) , *(L*^[*B*]^, *M* ^[*B*]^) , (*L*^[*V*]^, *M* ^[*V*]^).

Because DECODE’s clusters are ordered such that *p*_1_ *>* · · · *> p*_*H*_ , the cellular prevalence (***task 1A***) corresponds to the truncal cluster’s cancer cell fraction (CCF) *c*_1_ = 2 · *p*_1_. The prediction for subclone count (***task 1B***) is *H*. To score the inference of subclonal cellular prevalences (***task 1C***), DREAM requires the CCF and mutation count for each subclone. Similar to ***task 1A***, DECODE predicted the subclonal CCFs to be 2 · *p*_1_, … , 2 · *p*_*H*_. As the inference A subset has the lowest variant read count threshold (*L*^[*A*]^*< L*^[*B*]^, *L*^[*V*]^), we assumed that its deconvoluted SFS is the closest in shape to the unfiltered data, therefore the subclonal mutation counts follow ratio 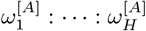.

DREAM’s scores for mutational assignment (***task 2A-B***) concern all mutations in the sample, and not limited to only diploid or heterozygous regions. Therefore, we first implemented GRITIC [Baker et al., 2024] to find the mutation multiplicities. For mutation *i* with observed VAF *f*_*i*_, multiplicity *m*_*i*_ and total copy number *c*_*i*_ in a sample with purity *ρ*, we defined its normalized VAF to be the expected VAF if the mutation were diploid [Tarabichi et al., 2021]:

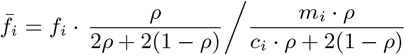

We then found the bin *b*_*i*_ that contained 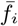. Because the SFS from inference A is closest in shape to the raw SFS, we expected the unfiltered mutation counts for different clusters in this bin to follow the ratio E(*S*^[*A*]^(*b*_*i*_) | *θ*_1 *H*_) : · · · : E (*S*^[*A*]^(*b*_*i*_) *θ*_*H*|*H*_). Therefore, the probability that mutation *i* to belong to cluster *h* is

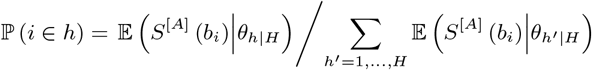

To determine the mutation’s hard assignment (***task 2A***), we attributed it to the cluster with the highest probability. The co-clustering matrix for ***task 2B*** consisted of the probabilities that each pair of mutations *i* and *j* belong to the same cluster:

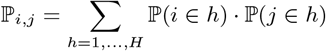

for *i* ≠ *j* and ℙ_*i,j*_ = 1 for *i* =*j*.

DECODE’s predictions for ***tasks 1A-2B*** were scored against ground truth using the evaluation scripts provided by the DREAM Challenge [Salcedo et al., 2020]. We then compared DECODE’s raw scores against 31 existing methods and five reference algorithms previously analyzed by DREAM’s authors [Salcedo et al., 2025] (Fig. 2 and Supplementary Figs. 1-3). To rank DECODE’s performance against the originally benchmarked algorithms, we computed the normalized score for each method in each task:

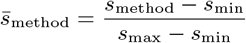

where *s*_method_, *s*_max_ and *s*_min_ are the method’s average raw score, and best and worst average scores among all software, respectively.

#### B.2 Comparison of computational costs

We compared the computational runtime between DECODE, PhyloWGS [Deshwar et al., 2015] and SciClone [Miller et al., 2014] across the 29 synthetic tumors included in our DREAM Challenge study. All methods were executed on the same hardware platform, a Mac mini (2023) with an Apple M2 Pro chip. Each package was executed with default settings. Any run that did not complete within 10^6^seconds (≈ 28 hours) was reported as incomplete.

### B.3 Detection and characterization of neutral tail

The ICGA-TCGA DREAM Challenge was not designed to validate the identification and profiling of the neutral tail component. Therefore, we performed a second benchmark study to quantify DECODE’s performance in detecting and characterizing the tail in the SFS.

We sampled each simulation’s purity *ρ*, total observed mutation count *m* and coverage distribution *φ* randomly and independently from real cancer samples in The Cancer Genome Atlas (TCGA, more details below) with mean coverage ≥ 30×. Each simulation further assumed *H* clones with hierarchy (*σ*_1_, … , *σ*_*H*_), where *σ*_*h*_ ∈ {0, … , *h*− 1} denotes the clonal parent of subclone *h* and 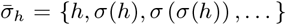 details its clonal lineage. Clone 1 was assumed to be truncal, hence *σ*_1_ = 0 and *σ*_2_, … , *σ*_*H*_ ≥ 1. The mutations in subclone *h*’s MRCA constituted a fraction 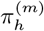 of the observed data (Fig. 1A), and the tail accounted for the remaining fraction 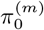. We assumed the sample consists of *n*_cancer_ = 10^5^cancer cells, a fraction 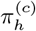 of which belongs to subclone *h*. The total cell count is then *n*_total_ = *n*_cancer_*/ρ*, contaminated by *n*_total_ − *n*_cancer_ normal cells, therefore there are 2 · *n*_total_ allele copies at any genomic site.

Each mutation was located at VAF *f* = *k/* (2 · *n*_total_) in the true neutral tail with probability 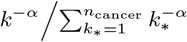, where *k* = 1, … , *n*_cancer_ is the number of cells (and hence allele copies) where the mutation is present, and *α* is the tail power. The mutation was sequenced with *r* reads with probability *φ*(*r*), then its variant read count was sampled from *z* ∼ Binomial(*r, f*). We filtered out mutations with *z* ≤ 2, mimicking typical variant-calling pipelines that exclude mutations supported by little evidence. Using this scheme, the simulator yielded *m* · *π*^(*m*)^tail mutations in the observed dataset. The *m*· *π*^(*m*)^observed mutations in each cluster *h* were simulated similarly with true VAF *f* = *n*_*h*_/(2 · *n*_total_) , where 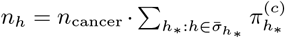 is the number of cells belonging to clone *h* and its subclones. The simulated mutational read count table was then analyzed by DECODE and MOBSTER [Caravagna et al., 2020], currently the only alternative mutation clustering algorithm that supports tail detection, with default settings for both methods.

The benchmark consisted of simulations with subclone count *H* = 1 (with hierarchy *σ*_1_ = 0), *H* = 2 (with (*σ*_1_, *σ*_2_) = (0, 1)) and *H* = 3 (with (*σ*_1_, *σ*_2_, *σ*_3_) = (0, 1, 1)), resulting in three simulation batches. Each batch consisted of 1800 simulated samples with tail (T-type) and 200 samples consisting of only mutation clusters without any tail mutations (NT-type). For each T-type simulation, tail power *α* was sampled from Uniform(0.9, 2.5), and both subclonal cell proportions 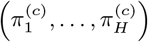 and mutation proportions 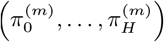 were sampled from Dirichlet(1, … , 1), conditioned on 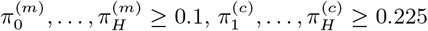 and min 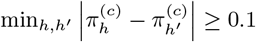. For NT-type simulations, cell proportions 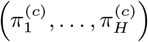 and mutation proportions 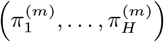 were similarly simulated and conditioned.

We evaluated decomposition results from DECODE and MOBSTER based on their detection and characterization of the tail component. The results were classified as false positive (FP) if they indicated the existence of a tail in NT-type simulations, and false negative (FN) if they failed to detect the tail in T-type samples. True-positive results were further evaluated based on the accuracy of the inferred tail power and tail mutation proportion, compared to the simulations’ ground-truth values of *α* and 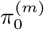. Both predictions were readily available from MOBSTER, as was the inferred tail power from DECODE. However, DECODE applied mutation filters before decomposing the SFS (Fig. 1B), hence its mutation counts were not directly comparable to those in the simulated data. DECODE’s decomposition of the *inference A* subsample containing *H* clusters can be represented as 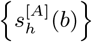, where 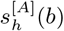 is the expected mutation count from compartment *h* in bin *b*. The corrected tail mutation proportion from DECODE is then:

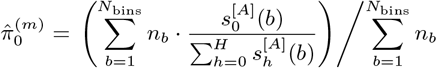

where *n*_*b*_ is the number of observed mutations in bin *b*.

We compared MOBSTER’s and DECODE’s accuracy in inferring tail parameter 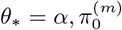 using root mean square error (RMSE):

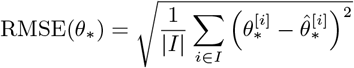

where 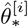 is the prediction compared to the true value 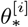 in simulation *i*, restricted to the subset *I* of simulations where the inferences of each method were true-positive.

We considered five confounding factors that may cause DECODE and MOBSTER’s results to be inaccurate: sample purity, coverage depth, mutation count, error in estimating cluster count, and the entropy caused by the overlap between the tail and mutation clusters. The cluster count error was defined as the inferred cluster count from each method minus the true cluster count. To quantify the tail entropy, let *n*_*h*_(*b*) be the number of mutations in bin *b* that belong to compartment *h* in the ground truth. We defined *p*_tail_(*b*) = *n*_0_(*b*)*/* _*h*_ *n*_*h*_(*b*) and *p*_clusters_(*b*) = 1 −*p*_tail_(*b*) to be the proportions of mutations in bin *b* that belong in the tail or clusters, respectively. The tail entropy was then based on Shannon’s diversity index:

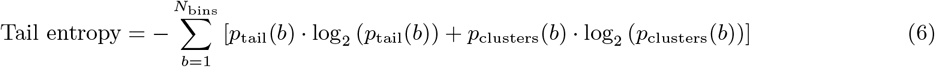

using the convention that 0 · log_2_(0) = 0. Higher tail entropy indicates greater overlap between the tail and mutation clusters, rendering the tail power and mutation count less identifiable (Supplementary Figs. 4-6).

### C Analysis of paired AML samples at diagnosis and relapse

Paired acute myeloid leukemia (AML) samples from 11 patients, collected at diagnosis and relapse, together with matched control samples described in [Shlush et al., 2017], were downloaded in BAM format from the EGA repository (ID: EGAD00001003234). The reads had previously been aligned with the hg19 reference genome using NovoAlign. All samples were additionally processed using Picard MarkDuplicates and GATK BaseRecalibrator (v4.1.1.0) [DePristo et al., 2011]. The average effective sequencing coverage was approximately 59×. Somatic variants were identified using GATK MuTect2 (v4.1.1.2) and filtered with FilterMutectCalls, incorporating contamination estimates from CalculateContamination and orientation bias statistics from LearnReadOrientationModel. Variants from matched diagnosis and relapse samples were combined using GATK CombineVariants, and variants failing filters in either sample were excluded to avoid inconsistencies caused by independent filtering. Retained variants were annotated using Variant Effect Predictor (VEP v97) [McLaren et al., 2016].

Copy number alterations were identified using GATK (v4.1.1.2) tools [DePristo et al., 2011] with the following tools and non-default parameters: PreprocessIntervals, CollectReadCounts, CreateReadCountPanelOfNormals (minimum-interval-median-percentile: 5.0), DenoiseReadCounts, CollectAllelicCounts, ModelSegments (number-of-smoothing-iterations-per-fit: 1, kernel-variance-copy-ratio: 0.8, kernel-variance-allele-fraction: 0.8, smoothing-credible-interval-threshold-allele-fraction: 10, smoothing-credible-interval-threshold-copy-ratio: 10), and CallCopyRatioSegments. These parameters were adjusted according to the authors’ recommendations for datasets exhibiting excessive numbers of CNV segments caused by noisy segment-specific read counts, as assessed using the GATK visualization tools PlotDenoisedCopyRatios and PlotModeledSegments.

DECODE was later used to analyze the paired samples at diagnosis and relapse (*n* = 11 patients). To remove samples with extremely low purity, we computed %_VAF*>*0.2_ for each patient, the percentage of diploid mutations with VAF *>* 0.2 at diagnosis or relapse, and excluded those with %_VAF*>*0.2_ < 5%. This eliminated Patient 9 (%_VAF*>*0.2_ = 2.4%) and retained ten patients (%_VAF*>*0.2_ = 12.5 − 60.6%).

For each patient, we implemented three mutational decomposition pipelines: (a) multi-sample SciClone [Miller et al., 2014] using paired data at diagnosis and relapse, (b) MOBSTER [Caravagna et al., 2020] and (c) DECODE, applied individually for each sample. The mutational data analyzed by all pipelines were filtered such that (1) multi-allelic loci were removed, (2) only diploid SNVs were retained, and (3) mutations with VAF *>* 0.75 were removed, as they likely occurred on regions with loss of heterozygosity (LOH), given that the copy number is 2. For multi-sample SciClone, mutations found in only one sample were given VAF = 0 in the other sample. By default, SciClone only analyzed mutations sequenced at depth ≥ 100× (minimumDepth = 100), which would remove 89 − 99% of mutations in each sample. Therefore, we applied SciClone on paired data with minimumDepth = 25 and default values for other hyper-parameters. MOBSTER and DECODE were conducted on single samples with default settings.

To illustrate the joint SFS to be expected under our proposed evolution model in Fig. 3I, we created synthetic data using TEMULATOR [Heide, 2022]. The simulation was configured to contain only one clone starting at time *t* = 0, evolving with birth rate 1.5, death rate 0.2 and mutation rate 200. The synthetic diagnosis and relapse samples were taken at time *t* = 7 and *t* = 10, respectively, such that each sample contained 500 cells. We assumed tumor purity = 1 in both simulated samples. Mutations were labeled “clonal” if their CCF ≥ 90% and “tail” otherwise. For each mutation *i* with CCF *c*_*i*_ in each sample, we first sampled its total read count *r*_*i*_ from the corresponding empirical coverage distribution from Patient 1, then generated its alternative read count as *z*_*i*_ ∼ Binomial(*r*_*i*_, *c*_*i*_*/*2). The joint and marginal SFS were then tallied from mutations with *z*_*i*_ ≥ 3.

To perform the dN/dS analysis, mutations from diagnosis and relapse were combined for each patient. The patient’s “cluster mutations” and “tail mutations” consisted of variants either classified accordingly in both samples, or assigned accordingly in one sample and undetected in the other. We then implemented R package dndscv [Martincorena et al., 2017] to compute the dN/dS ratio for each mutation category and patient.

### D Analysis of primary tumors in TCGA

We applied DECODE to tumor samples from The Cancer Genome Atlas (TCGA) [Weinstein et al., 2013]. The cohort consisted of 10,106 samples for which both mutation calls from MuTect2 [Benjamin et al., 2019] and allele-specific copy number profiles from ASCAT3 [Van Loo et al., 2010] were available, across 8,033 patients and 32 cancer types. Our analysis was limited to 4,666 samples with mean coverage ≥ 30× that were predominantly diploid, i.e., with ≥ 60% mutations residing on diploid regions (major CN = minor CN = 1) and ≥ 50 diploid mutations in total.

We examined DECODE’s consistency by analyzing 57 pairs of biological replicates across 49 patients in TCGA. DECODE and SciClone [Miller et al., 2014] were implemented to cluster mutations in individual samples, restricted to diploid mutations as required by both algorithms. Similar to the AML study, SciClone was applied with minimumDepth = 25, and default values for other hyper-parameters. DECODE was applied in the default configuration, i.e., the best fits with and without tail were compared to determine the parsimonious deconvolutions. Although DECODE finished all analyses, SciClone did not complete for two patients (TCGA-DJ-A1QF and TCGA-AZ-4315) within 1 day; these patients were thus excluded in SciClone’s results. We compared the assignments of mutations to SFS compartments (i.e., either the tail or specific clusters) across biological replicates from each algorithm to evaluate its consistency. For each patient, we included mutations that were (i) present in both replicates, (ii) located in diploid regions, and (iii) assigned to specific compartments in each replicate. The third condition excluded mutations unassigned by SciClone due to their locations in the SFS being where multiple subclones overlapped. The adjusted Rand index (ARI) for the biological replicate pair is then

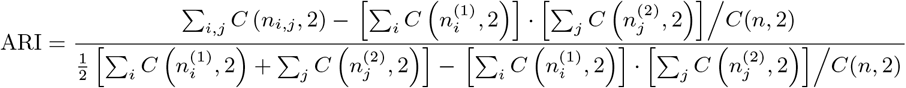

where *n* is the total mutation count, 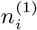 is the number of mutations in compartment *i* in replicate 1, 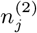 is the number of mutations in compartment *j* in replicate 2, *n*_*i,j*_ is the number of mutations assigned to compartment *i* in replicate 1 and compartment *j* in replicate 2, and 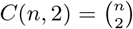. ARI scores for pairs of biological replicates were computed with R package mclust [Scrucca et al., 2023], then compared between DECODE and SciClone.

We also evaluated the consistency in DECODE-inferred tail powers. Among the 49 patients with multi-region WGS data, DECODE detected the tail in more than one region in 29 patients, resulting in 35 pairs of biological replicates where tail powers were assigned for both samples. For each pair of same-patient samples (*i, j*) with tail powers *α*_*i*_ and *α*_*j*_ , we defined the pairwise *α* difference as *d*(*α*_*i*_, *α*_*j*_) = |*α*_*i*_ − *α*_*j*_ |. We then performed the permutation test, where the observed median same-patient *α* difference was compared to the medians of *d*(*α*_*i*_, *α*_*j*_) where *i* and *j* were randomly permuted among samples from all patients. The reported p-value reflected the proportion of 10,000 permutations that resulted in lower medians than the true median same-patient difference.

We next examined DECODE’s results for the whole diploid TCGA dataset. To evaluate the relationship between patient characteristics and outcome on one hand and DECODE-inferred tail power and clonality at diagnosis on the other, we restricted the analysis to one sample per patient, selecting the sample with highest coverage when biological or technical replicates were available. Our analysis then consisted of *n* = 4,204 unique tumors across 32 cancer types. For each sample, the input for DECODE likewise consisted only of diploid mutations.

We compared sample purities predicted from DECODE against the values inferred previously from ASCAT3. As in the DREAM Challenge study, the purities can be estimated from truncal cluster VAFs. However, inaccurate copy number inference can produce spurious subclones with VAF ≫ 0.5 (i.e., CCF ≫ 1), for instance due to genomic regions with (major CN, minor CN) = (2, 0) being miscalled as (1, 1). Therefore, DECODE’s inferred sample purity was computed as 2 × max {*p*_*h*_ : *p*_*h*_ ≤ 0.55, *h* = 1, … , *H*}. On the other hand, ASCAT3 depends on the genomic regions with diverse copy number states to estimate purity. As a result, purity estimates were unreliable for samples with low aneuploidy (for these cases, ASCAT3 defaulted to purity = 1). We therefore restricted the comparison of purity estimates between DECODE and ASCAT3 to samples in which 10% of the genome or more was aneuploid (*n* = 2,333).

To standardize our comparison of tumor characteristics, DECODE deconvolutions and patient outcomes, we limited the analysis to de novo cancers (4,058 unique tumors across 32 types) and harmonized AJCC pathologic stage and tumor grade annotations into broader categories. AJCC stage annotations were mapped as follows: stages 0, 0a, 0is and IS were grouped as stage 0; stages I, IA, IB, IB1 as stage I; stages II, IIA, IIB and IIC as stage II; stages III, IIIA, IIIB and IIIC as stage III; and stages IV, IVA, IVB, IVC and X as stage IV; with missing or other annotations being grouped as unknown. Two cases warrant note: stage IS, observed in one TGCT patient (`TCGA-2G-AAF1`), was grouped into stage 0 following clinical staging convention, and stage X, observed in four BRCA patients, was grouped with stage IV. Tumor grade was primarily grouped into G1, G2, G3 and G4. Grades GB, observed in one OV patient (`TCGA-24-2038`), and GX, observed in 23 patients across CESC, ESCA, HNSC, KIRC, PAAD, and STAD cancer types, were treated as unknown. For BLCA, where tumors were annotated as low grade or high grade, these categories were retained as reported; high grade was also observed in 2 of 294 UCEC patients and remained as such.

We examined which patient subsets exhibited an association between DECODE-inferred clonality and treatment outcome. We stratified TCGA primary cancers into cohorts by (i) cancer type, (ii) cancer type and AJCC stage, or (iii) cancer type and tumor grade. Samples in each cohort were partitioned into two subgroups by comparing their DECODE-derived cluster counts to the cohort median. Survival analysis was performed only for cohorts in which (a) both subgroups contained ≥ 20 patients, and (b) each subgroup included ≥ five death events. A total of 26 cohorts met these criteria: 15 were defined by cancer type alone, three by cancer type and AJCC stage, and eight by cancer type and tumor grade. For each cohort, the hazard ratio (HR) was computed from a Cox proportional hazards model with subgroup identity as the predictor of patient survival. Differences in survival between the two subgroups were evaluated using the log-rank test applied to Kaplan–Meier survival curves. The resulting p-values were adjusted for multiple testing using the Benjamini–Hochberg (BH) procedure, yielding BH-adjusted p-values *p*_adj_.

We investigated two TCGA cohorts where partitioning by median cluster count led to statistically significant differences in survival: LGG samples, and LGG grade 3 tumors, both with median cluster count = 1. We first compared tail powers between the partitioned subgroups across both cohorts, limited to samples where DECODE detected a tail (LGG: *n* = 301, LGG grade 3: *n* = 127). We then classified an LGG sample as IDH mutant if it contained one or more mutations that (i) affected IDH1 or IDH2, (ii) passed MuTect2 filtering, and (iii) had moderate or high predicted functional impact (LGG: *n* = 231, LGG grade 3: *n* = 87). We compared the proportion of samples where DECODE detected multiple subclones in each cohort, partitioned by LGG mutant and LGG wild-type, using two-sided Fisher’s exact test.

## Code availability

DECODE is available at https://github.com/dinhngockhanh/DECODE.

## Acknowledgments

YC, MK and KND were supported by the National Cancer Institute of the National Institutes of Health under Award Number R01CA310281. RJ, MKK and MK were supported by the Polish National Science Centre under Award Number UMO-2021/41/B/NZ2/04134. The content is solely the responsibility of the authors and does not necessarily represent the official views of the National Institutes of Health.

---

**Supplementary Figure 1:**
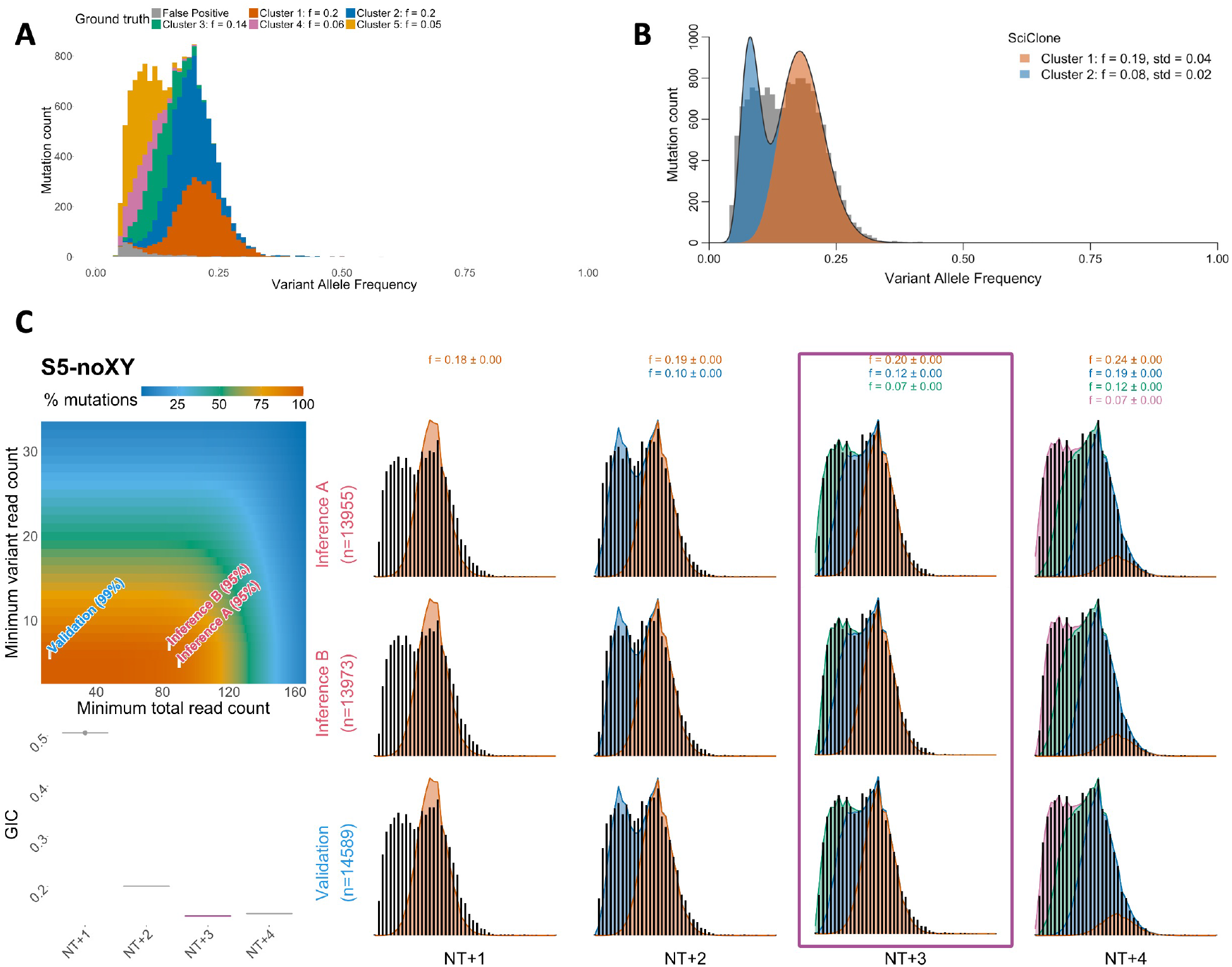
Nonexistent clonal cluster results in incorrectly inferred sample purity. **A-B**: Ground truth SFS (**A**) and SciClone’s decomposition (**B**) for diploid mutations in DREAM synthetic tumor S5. **C**: DECODE’s decomposition of tumor S5. Top left: heatmap of mutation retention frequency for different variant and total read count thresholds. Filtering thresholds for *inference A, inference B* and *validation* subsamples are highlighted. Right: Optimal decompositions given cluster count *H* under assumption of no tail (column NT+*H*). For each *H*, the expected observed SFS from the posterior distribution for parameters *θ*_*H*_ (shaded areas, colors denoting clusters) for each subsample is compared against the empirical SFS (black histogram). Means and standard deviations of cluster VAFs are reported on top. Bottom left: Comparison of Generalized Information Criteria (GIC) between different cluster count assumptions. The fit with lowest GIC (NT+3) is chosen.

**Supplementary Figure 2:**
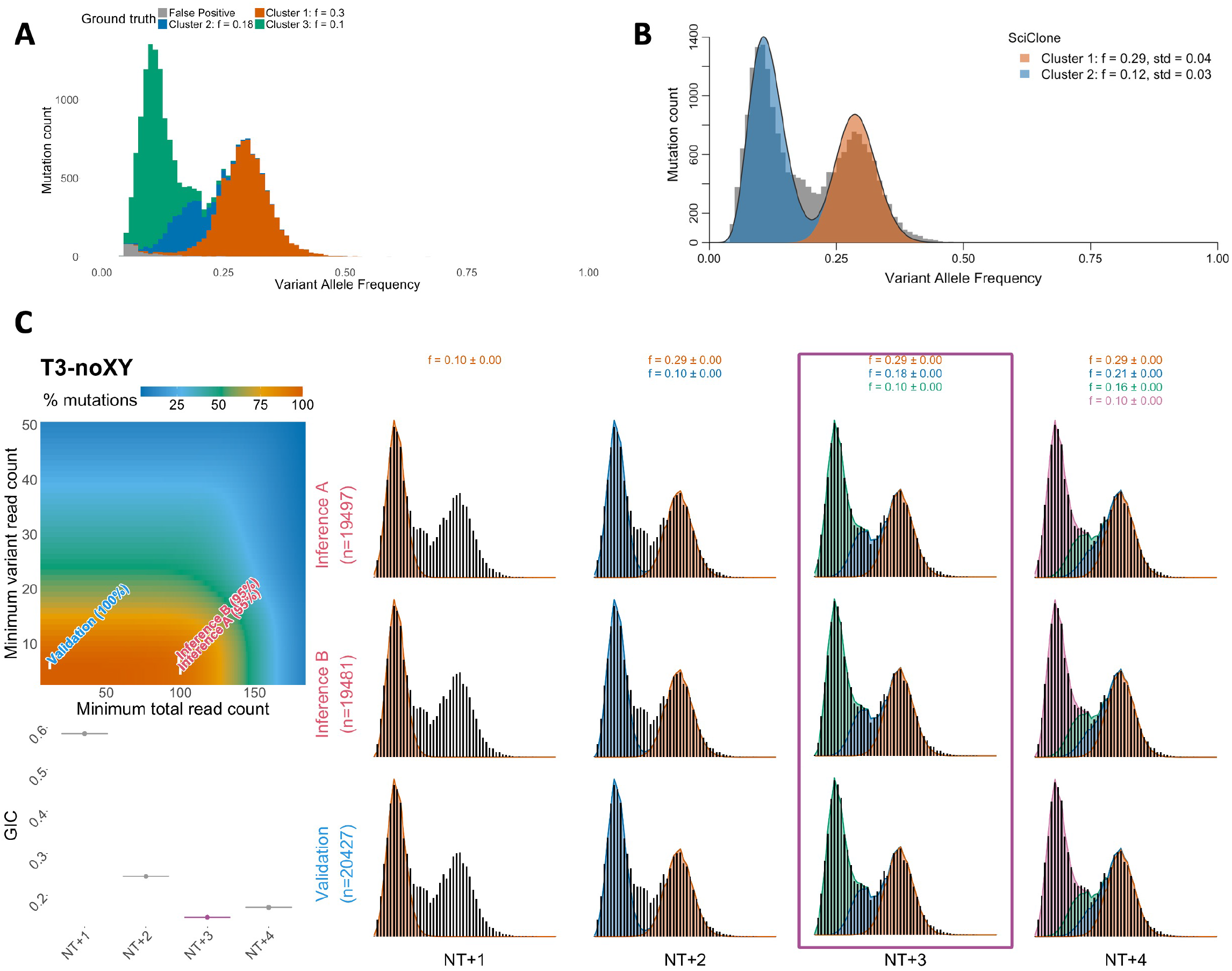
DECODE accurately identifies obscured clusters. **A-B**: Ground truth SFS (**A**) and SciClone’s decomposition (**B**) for diploid mutations in DREAM synthetic tumor T3. **C**: DECODE’s decomposition of tumor T3. Top left: heatmap of mutation retention frequency for different variant and total read count thresholds. Filtering thresholds for *inference A, inference B* and *validation* subsamples are highlighted. Right: Optimal decompositions given cluster count *H* under assumption of no tail (column NT+*H*). For each *H*, the expected observed SFS from the posterior distribution for parameters *θ*_*H*_ (shaded areas, colors denoting clusters) for each subsample is compared against the empirical SFS (black histogram). Means and standard deviations of cluster VAFs are reported on top. Bottom left: Comparison of Generalized Information Criteria (GIC) between different cluster count assumptions. The fit with lowest GIC (NT+3) is chosen.

**Supplementary Figure 3:**
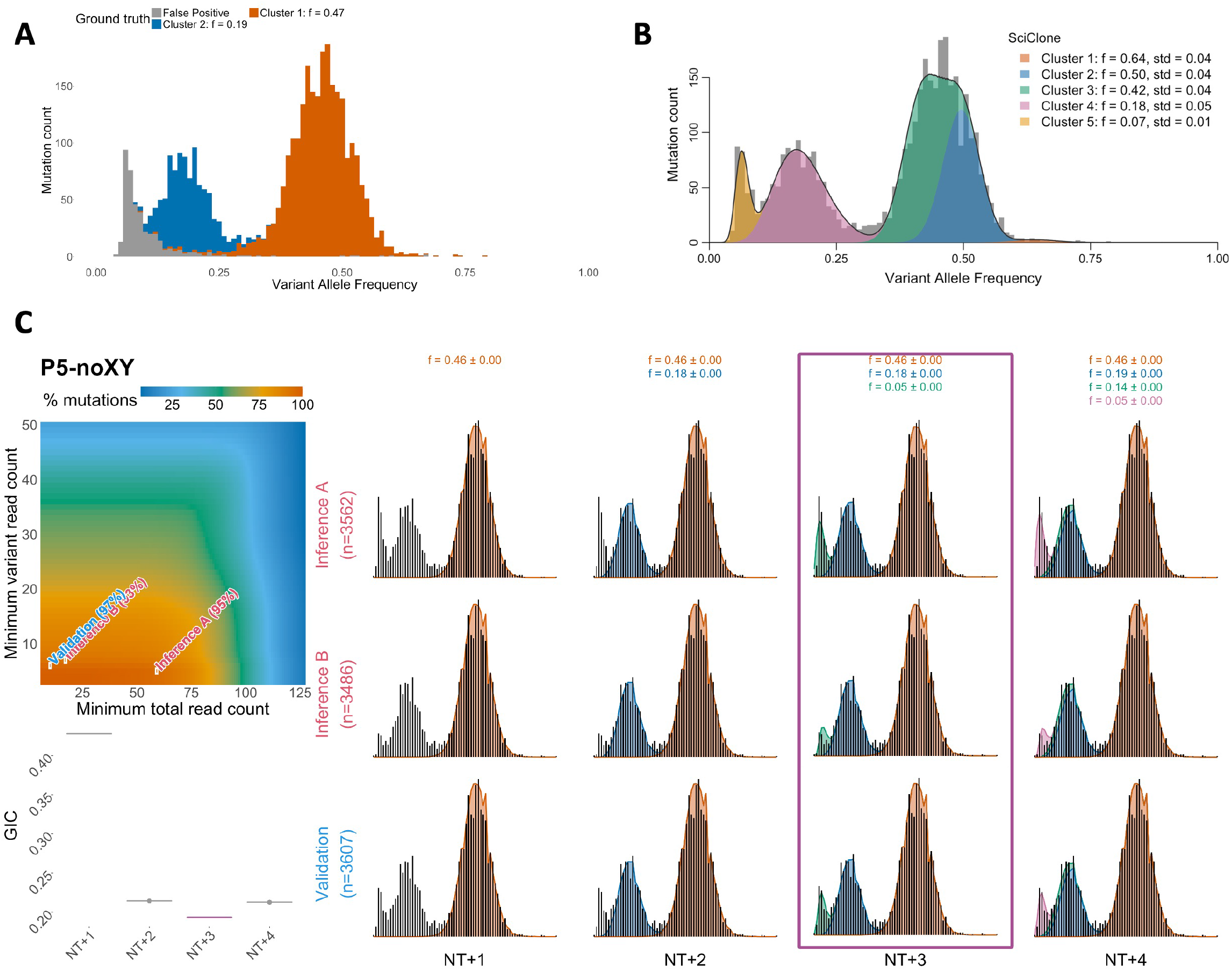
DECODE mitigates cluster overfitting. **A-B**: Ground truth SFS (**A**) and SciClone’s decomposition (**B**) for diploid mutations in DREAM synthetic tumor P5. **C**: DECODE’s decomposition of tumor P5. Top left: heatmap of mutation retention frequency for different variant and total read count thresholds. Filtering thresholds for *inference A, inference B* and *validation* subsamples are highlighted. Right: Optimal decompositions given cluster count *H* under assumption of no tail (column NT+*H*). For each *H*, the expected observed SFS from the posterior distribution for parameters *θ*_*H*_ (shaded areas, colors denoting clusters) for each subsample is compared against the empirical SFS (black histogram). Means and standard deviations of cluster VAFs are reported on top. Bottom left: Comparison of Generalized Information Criteria (GIC) between different cluster count assumptions. The fit with lowest GIC (NT+3) is chosen.

**Supplementary Figure 4:**
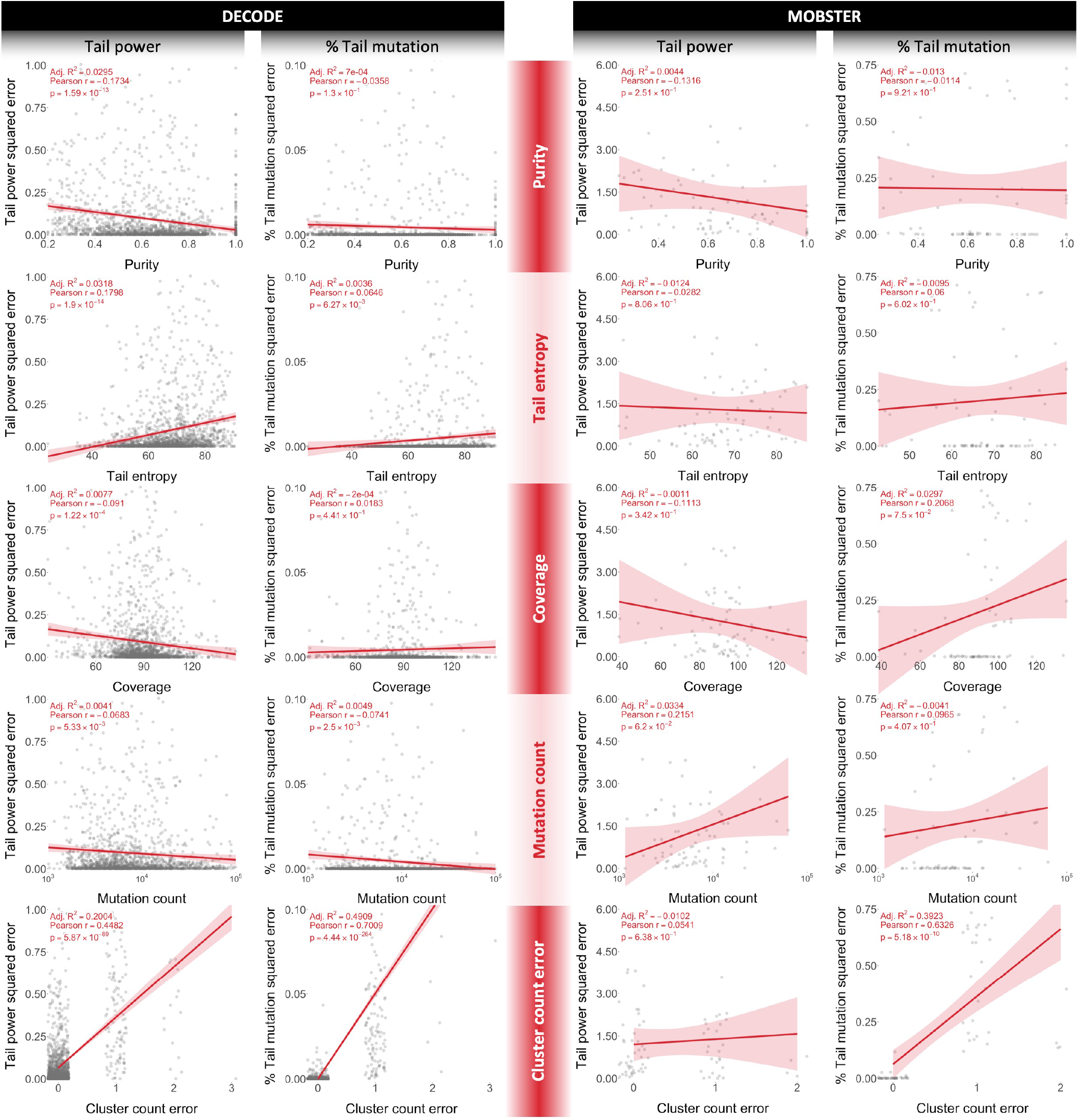
Analysis of DECODE’s and MOBSTER’s errors in estimating tail power and mutation proportion from synthetic samples with true cluster count *H* = 1. Plots display each method’s squared errors from individual simulations (dots) as a function of sample purity, tail entropy, coverage depth, observed mutation count, and error in estimating cluster count. Synthetic samples’ purity, coverage and mutation count are drawn randomly from TCGA samples. Tail entropy is calculated using Eq. (6). Cluster count error equals the inferred cluster count minus the true value. Red line indicates the linear regression fit, annotations report Pearson’s correlation coefficient, its p-value, and the adjusted *R*^2^.

**Supplementary Figure 5:**
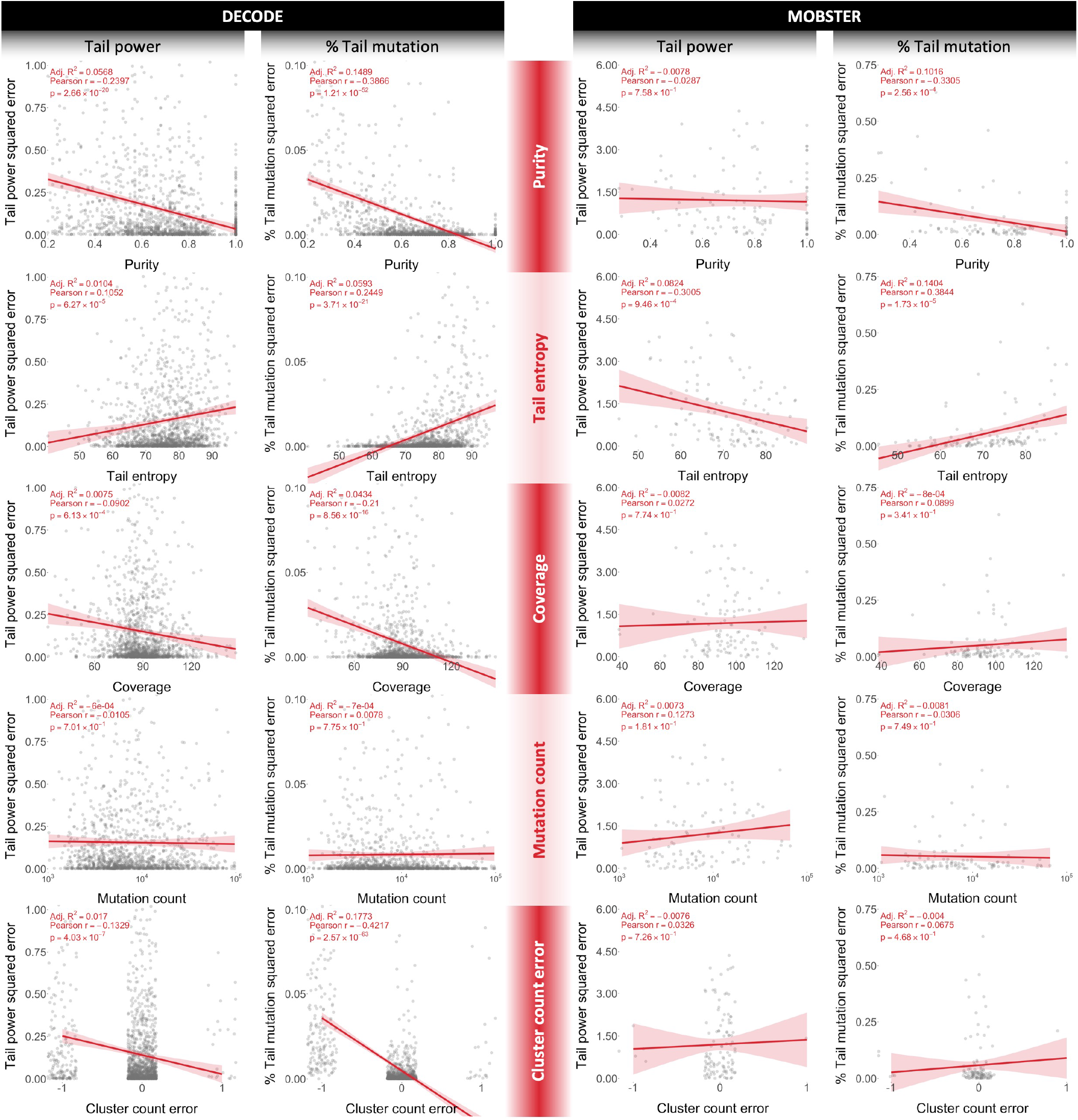
Analysis of DECODE’s and MOBSTER’s errors in estimating tail power and mutation proportion from synthetic samples with true cluster count *H* = 2. Plots display each method’s squared errors from individual simulations (dots) as a function of sample purity, tail entropy, coverage depth, observed mutation count, and error in estimating cluster count. Synthetic samples’ purity, coverage and mutation count are drawn randomly from TCGA samples. Tail entropy is calculated using Eq. (6). Cluster count error equals the inferred cluster count minus the true value. Red line indicates the linear regression fit, annotations report Pearson’s correlation coefficient, its p-value, and the adjusted *R*^2^.

**Supplementary Figure 6:**
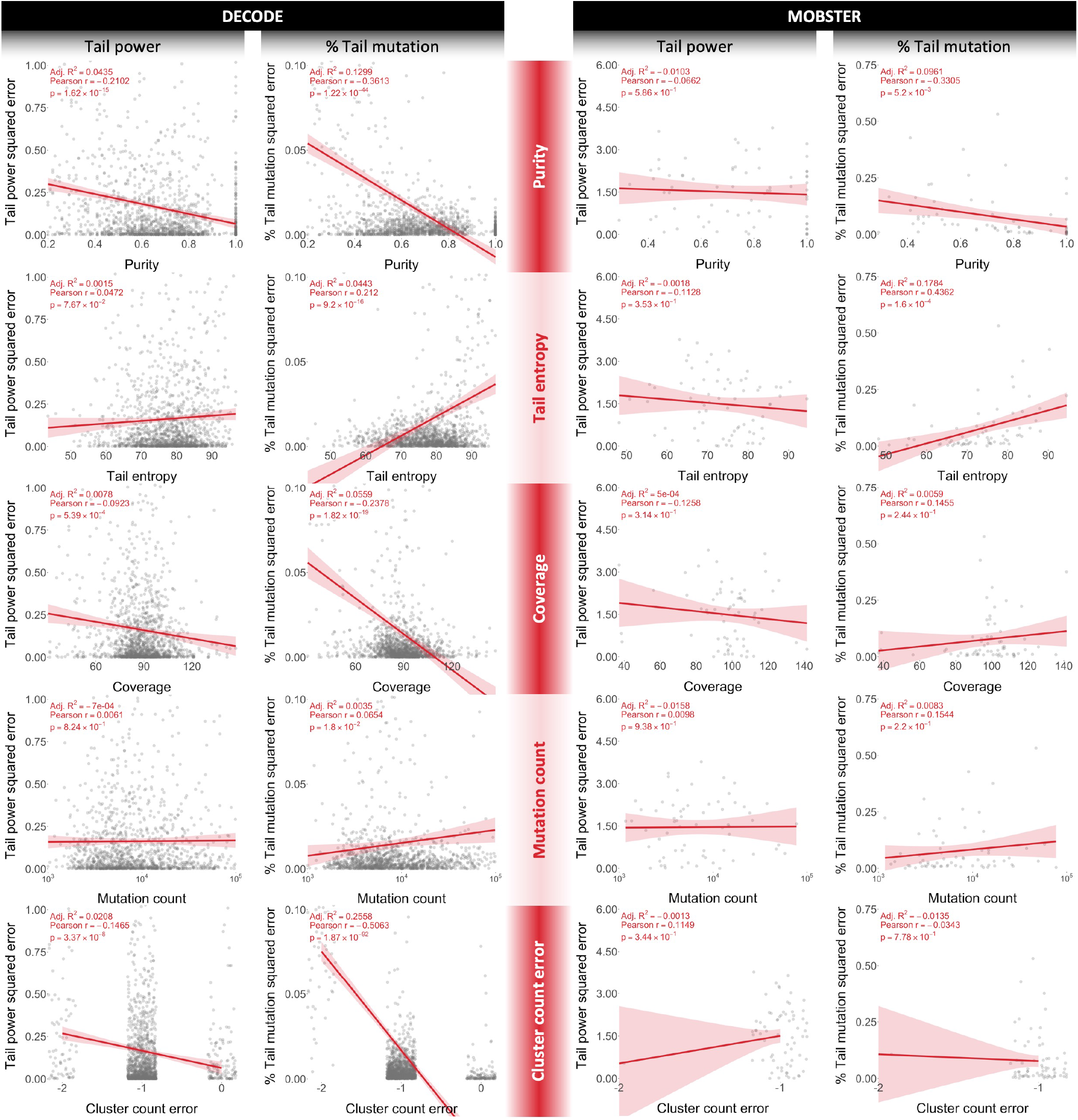
Analysis of DECODE’s and MOBSTER’s errors in estimating tail power and mutation proportion from synthetic samples with true cluster count *H* = 3. Plots display each method’s squared errors from individual simulations (dots) as a function of sample purity, tail entropy, coverage depth, observed mutation count, and error in estimating cluster count. Synthetic samples’ purity, coverage and mutation count are drawn randomly from TCGA samples. Tail entropy is calculated using Eq. (6). Cluster count error equals the inferred cluster count minus the true value. Red line indicates the linear regression fit, annotations report Pearson’s correlation coefficient, its p-value, and the adjusted *R*^2^.

**Supplementary Figure 7:**
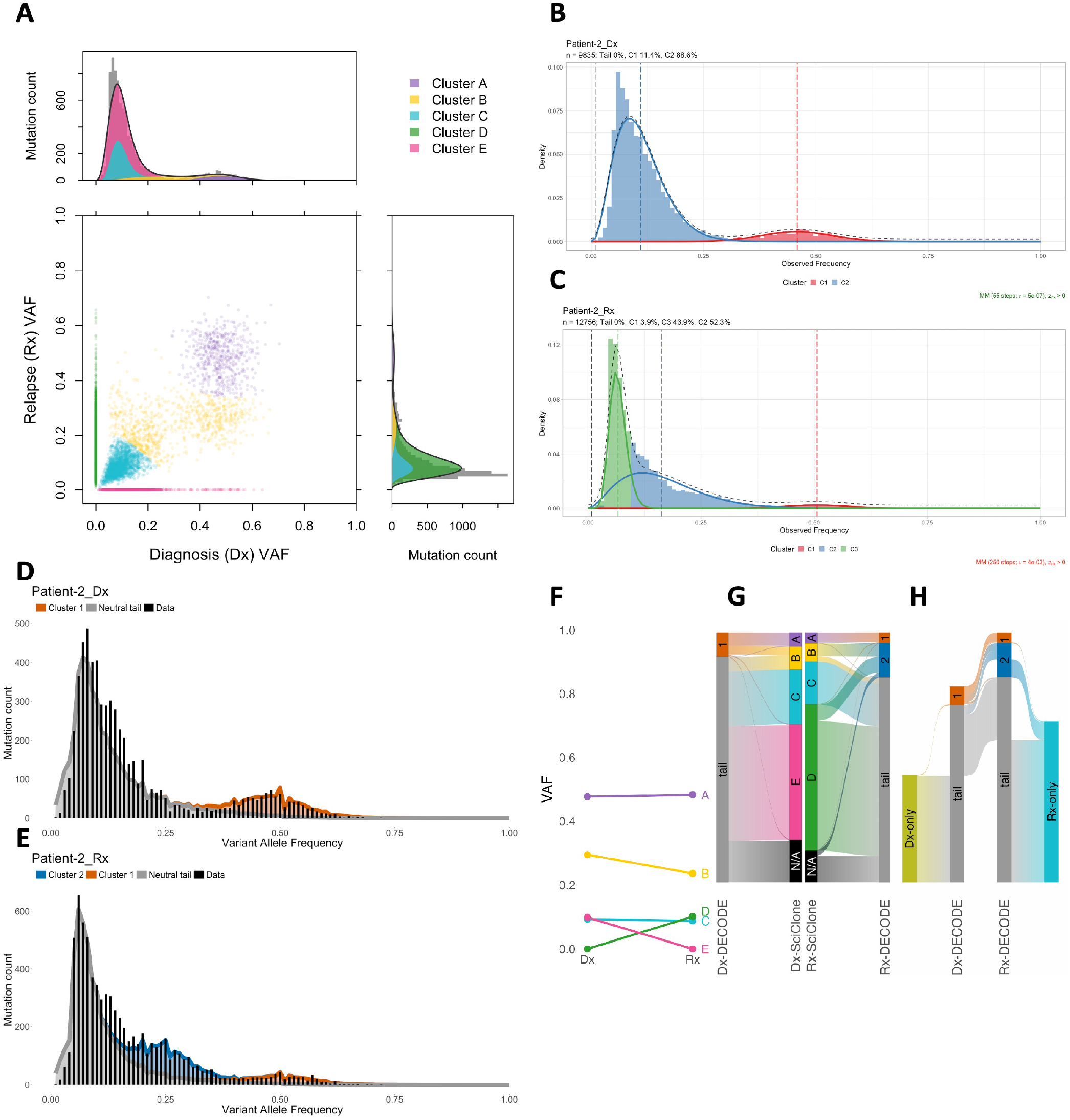
Mutation clustering for patient AML-2. **A**: SciClone’s joint clustering of mutations at diagnosis and relapse. **B-E**: Deconvolution of mutations using MOBSTER (**B, C**) and DECODE (**D, E**) for each sample at diagnosis (**B, D**) and relapse (**C, E**). **F**: Mean VAFs between diagnosis and relapse of subclones inferred from SciClone. **G**: Alluvial diagrams of subclonal assignments for mutations at diagnosis (left) and relapse (right) between DECODE and SciClone. Widths are proportional to mutation fractions. **H**: Alluvial diagram of DECODE subclonal assignments for mutations at diagnosis and relapse. Outer bars contain mutations unique to each sample.

**Supplementary Figure 8:**
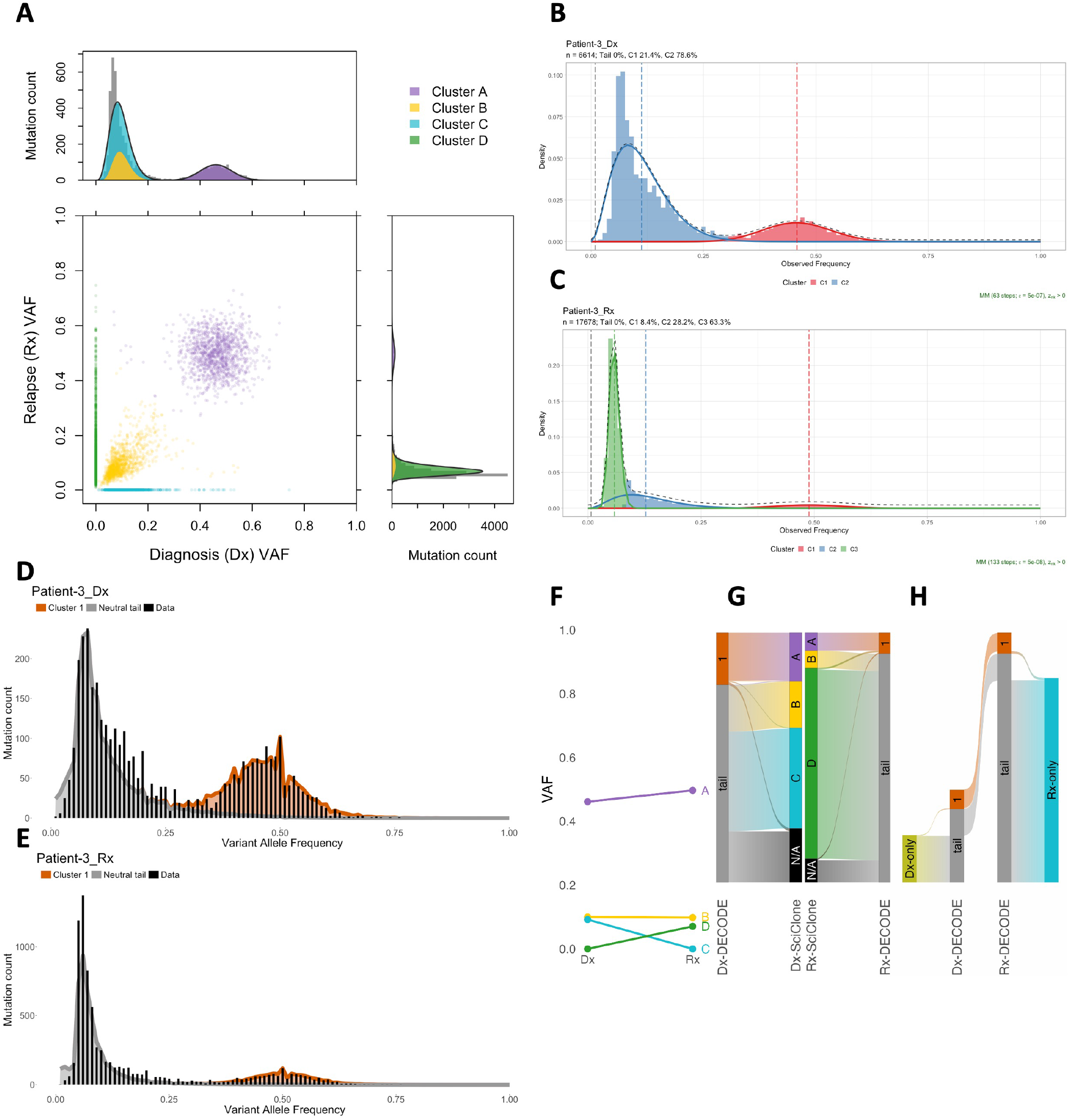
Mutation clustering for patient AML-3. **A**: SciClone’s joint clustering of mutations at diagnosis and relapse. **B-E**: Deconvolution of mutations using MOBSTER (**B, C**) and DECODE (**D, E**) for each sample at diagnosis (**B, D**) and relapse (**C, E**). **F**: Mean VAFs between diagnosis and relapse of subclones inferred from SciClone. **G**: Alluvial diagrams of subclonal assignments for mutations at diagnosis (left) and relapse (right) between DECODE and SciClone. Widths are proportional to mutation fractions. **H**: Alluvial diagram of DECODE subclonal assignments for mutations at diagnosis and relapse. Outer bars contain mutations unique to each sample.

**Supplementary Figure 9:**
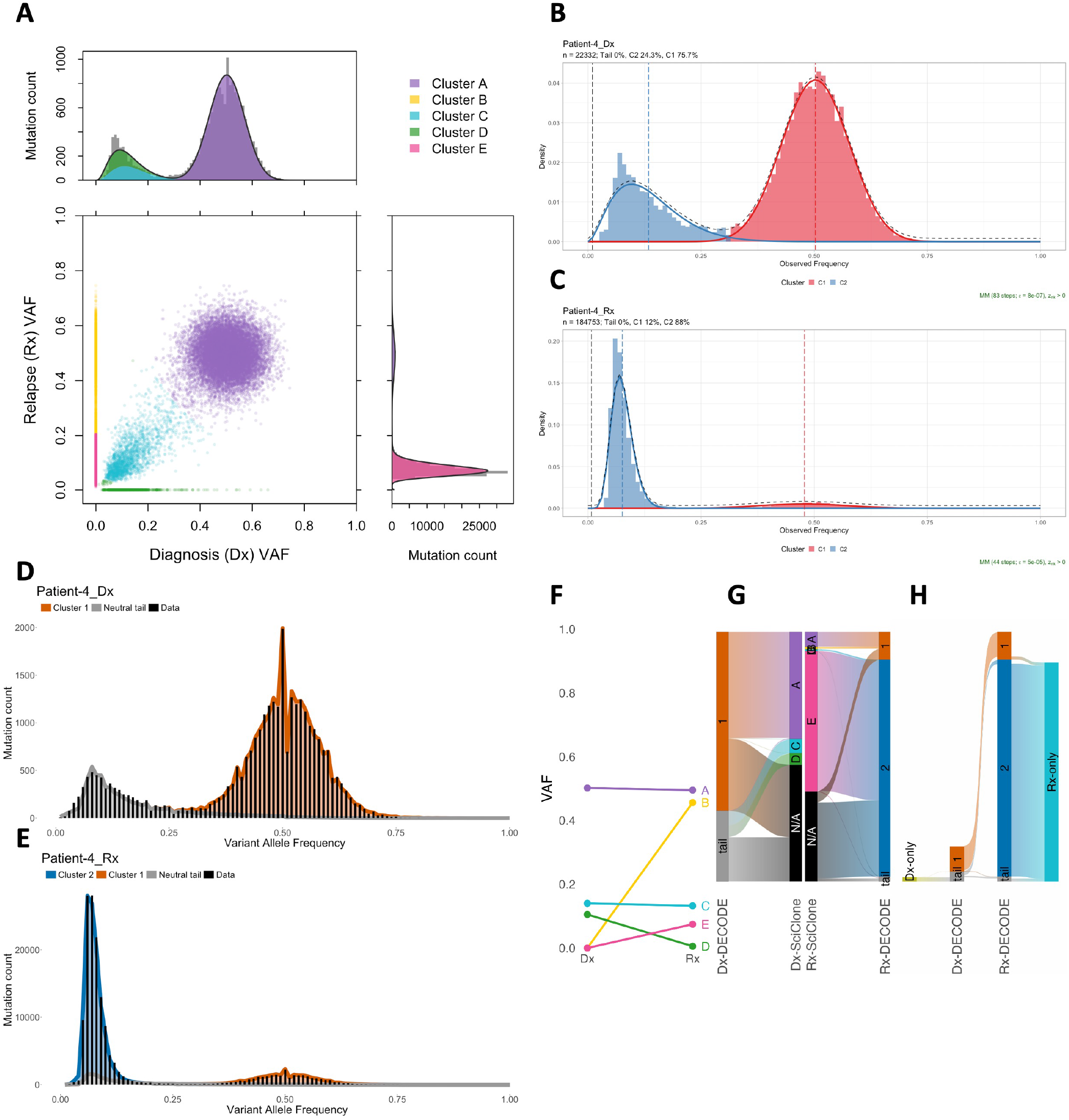
Mutation clustering for patient AML-4. **A**: SciClone’s joint clustering of mutations at diagnosis and relapse. **B-E**: Deconvolution of mutations using MOBSTER (**B, C**) and DECODE (**D, E**) for each sample at diagnosis (**B, D**) and relapse (**C, E**). **F**: Mean VAFs between diagnosis and relapse of subclones inferred from SciClone. **G**: Alluvial diagrams of subclonal assignments for mutations at diagnosis (left) and relapse (right) between DECODE and SciClone. Widths are proportional to mutation fractions. **H**: Alluvial diagram of DECODE subclonal assignments for mutations at diagnosis and relapse. Outer bars contain mutations unique to each sample.

**Supplementary Figure 10:**
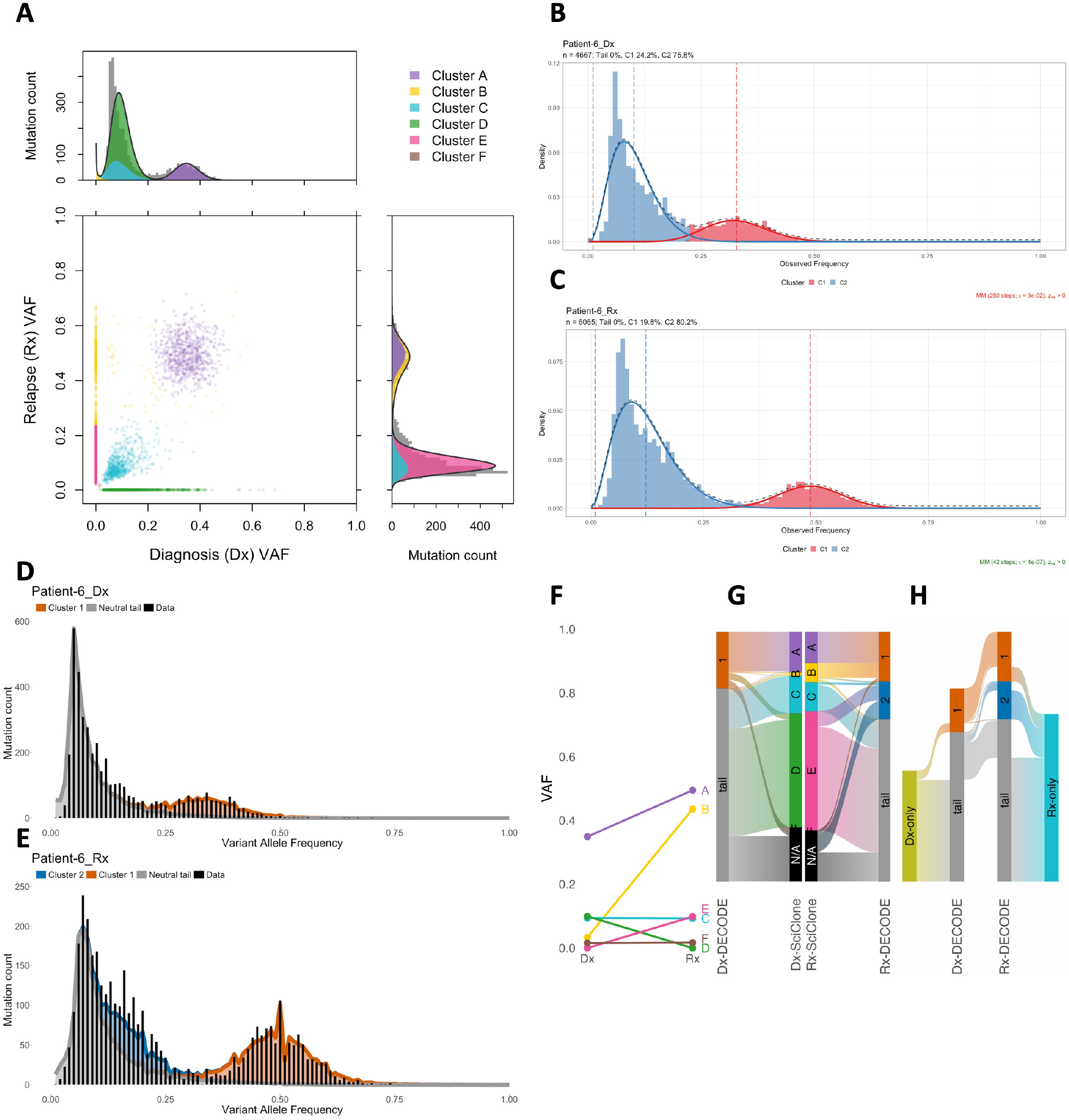
Mutation clustering for patient AML-6. **A**: SciClone’s joint clustering of mutations at diagnosis and relapse. **B-E**: Deconvolution of mutations using MOBSTER (**B, C**) and DECODE (**D, E**) for each sample at diagnosis (**B, D**) and relapse (**C, E**). **F**: Mean VAFs between diagnosis and relapse of subclones inferred from SciClone. **G**: Alluvial diagrams of subclonal assignments for mutations at diagnosis (left) and relapse (right) between DECODE and SciClone. Widths are proportional to mutation fractions. **H**: Alluvial diagram of DECODE subclonal assignments for mutations at diagnosis and relapse. Outer bars contain mutations unique to each sample.

**Supplementary Figure 11:**
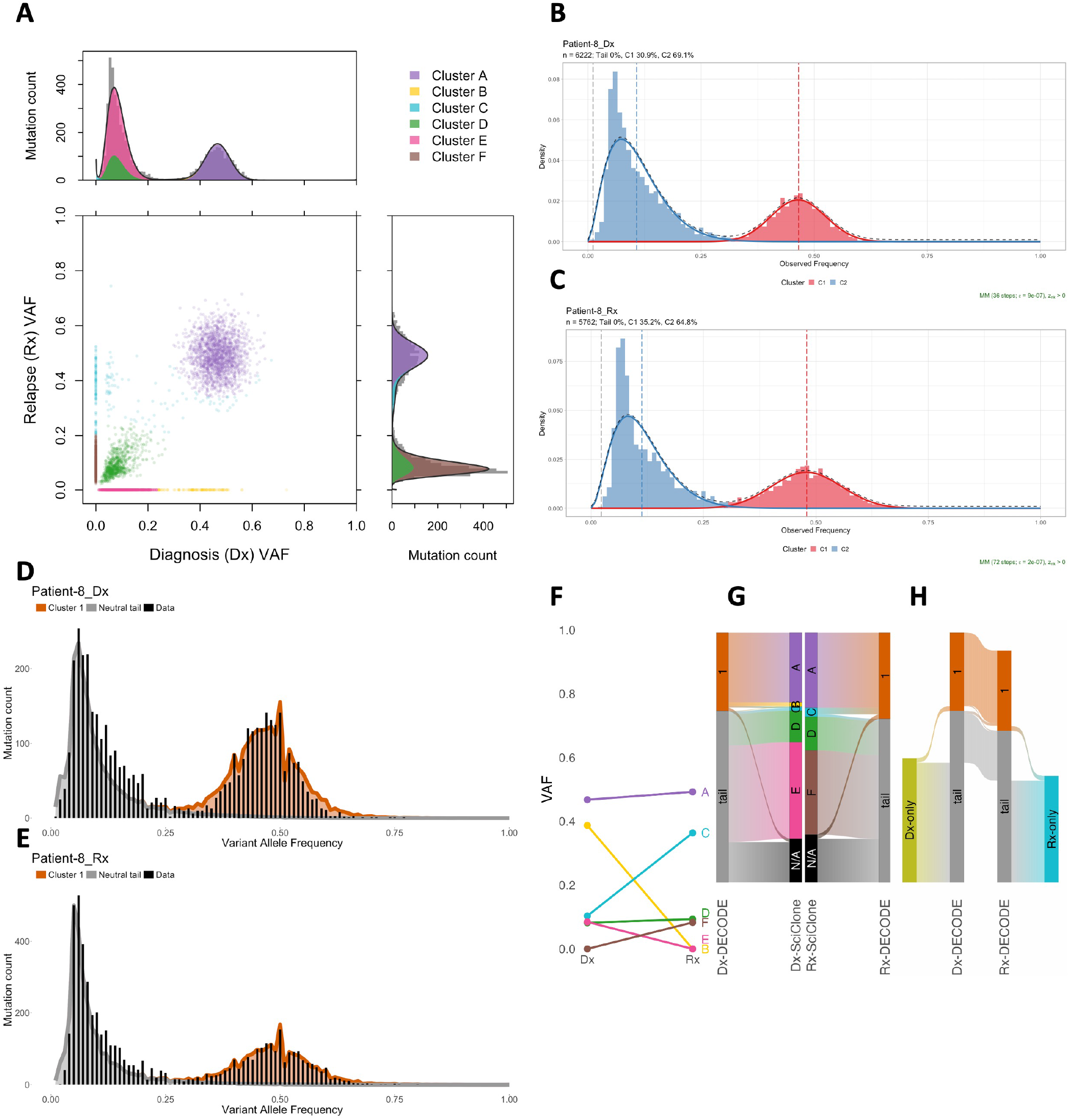
Mutation clustering for patient AML-8. **A**: SciClone’s joint clustering of mutations at diagnosis and relapse. **B-E**: Deconvolution of mutations using MOBSTER (**B, C**) and DECODE (**D, E**) for each sample at diagnosis (**B, D**) and relapse (**C, E**). **F**: Mean VAFs between diagnosis and relapse of subclones inferred from SciClone. **G**: Alluvial diagrams of subclonal assignments for mutations at diagnosis (left) and relapse (right) between DECODE and SciClone. Widths are proportional to mutation fractions. **H**: Alluvial diagram of DECODE subclonal assignments for mutations at diagnosis and relapse. Outer bars contain mutations unique to each sample.

**Supplementary Figure 12:**
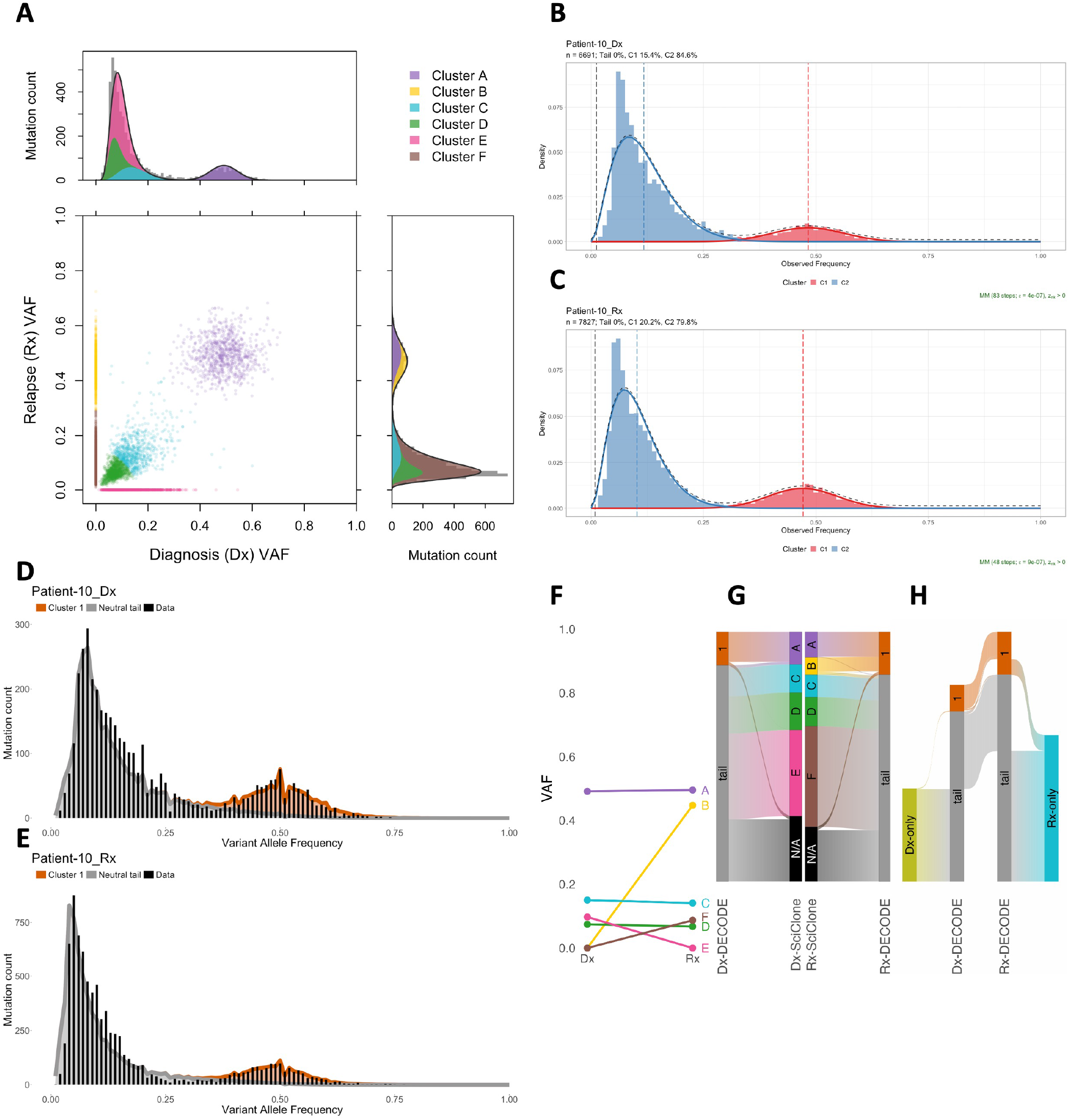
Mutation clustering for patient AML-10. **A**: SciClone’s joint clustering of mutations at diagnosis and relapse. **B-E**: Deconvolution of mutations using MOBSTER (**B, C**) and DECODE (**D, E**) for each sample at diagnosis (**B, D**) and relapse (**C, E**). **F**: Mean VAFs between diagnosis and relapse of subclones inferred from SciClone. **G**: Alluvial diagrams of subclonal assignments for mutations at diagnosis (left) and relapse (right) between DECODE and SciClone. Widths are proportional to mutation fractions. **H**: Alluvial diagram of DECODE subclonal assignments for mutations at diagnosis and relapse. Outer bars contain mutations unique to each sample.

**Supplementary Figure 13:**
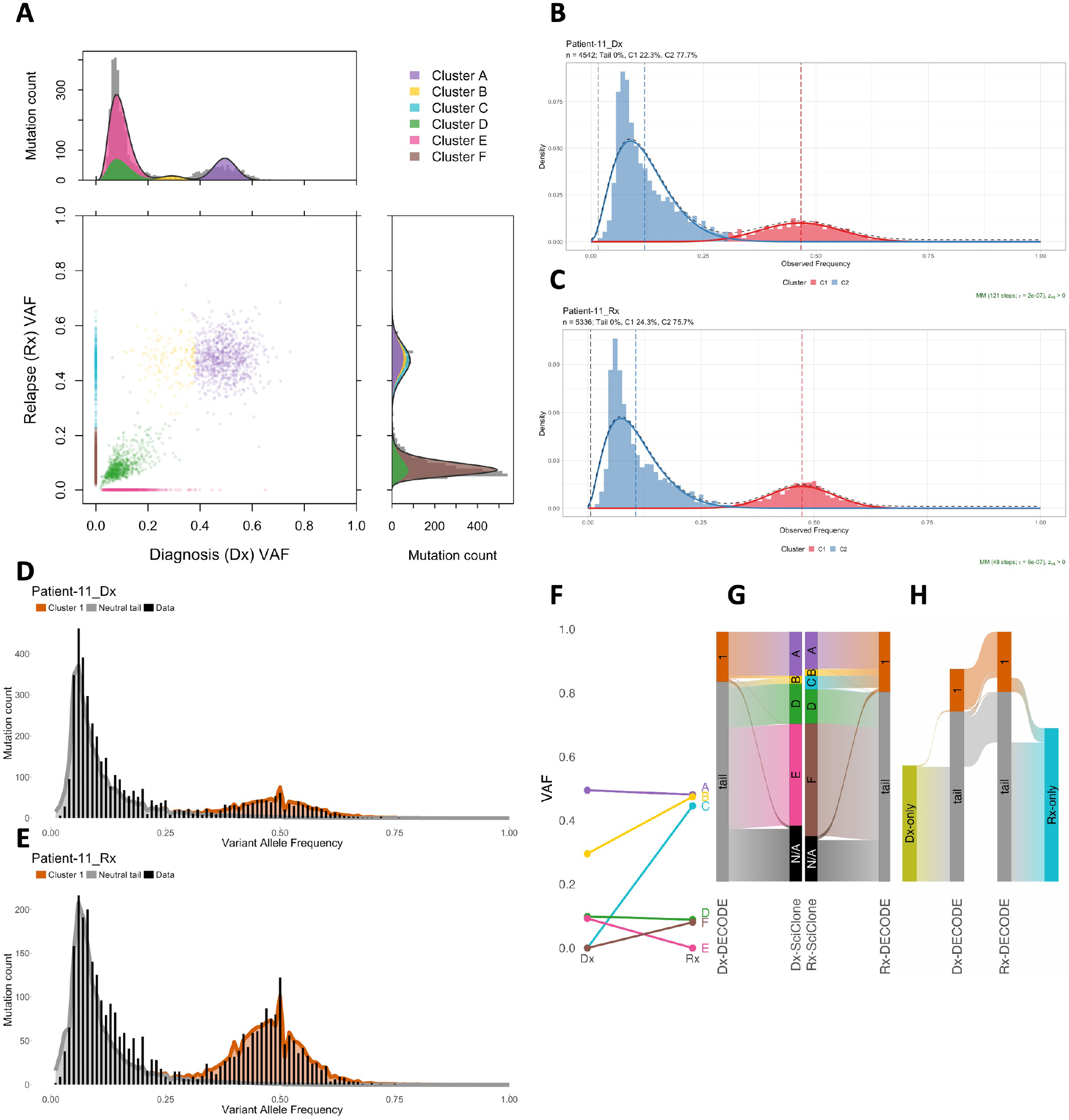
Mutation clustering for patient AML-11. **A**: SciClone’s joint clustering of mutations at diagnosis and relapse. **B-E**: Deconvolution of mutations using MOBSTER (**B, C**) and DECODE (**D, E**) for each sample at diagnosis (**B, D**) and relapse (**C, E**). **F**: Mean VAFs between diagnosis and relapse of subclones inferred from SciClone. **G**: Alluvial diagrams of subclonal assignments for mutations at diagnosis (left) and relapse (right) between DECODE and SciClone. Widths are proportional to mutation fractions. **H**: Alluvial diagram of DECODE subclonal assignments for mutations at diagnosis and relapse. Outer bars contain mutations unique to each sample.

**Supplementary Figure 14:**
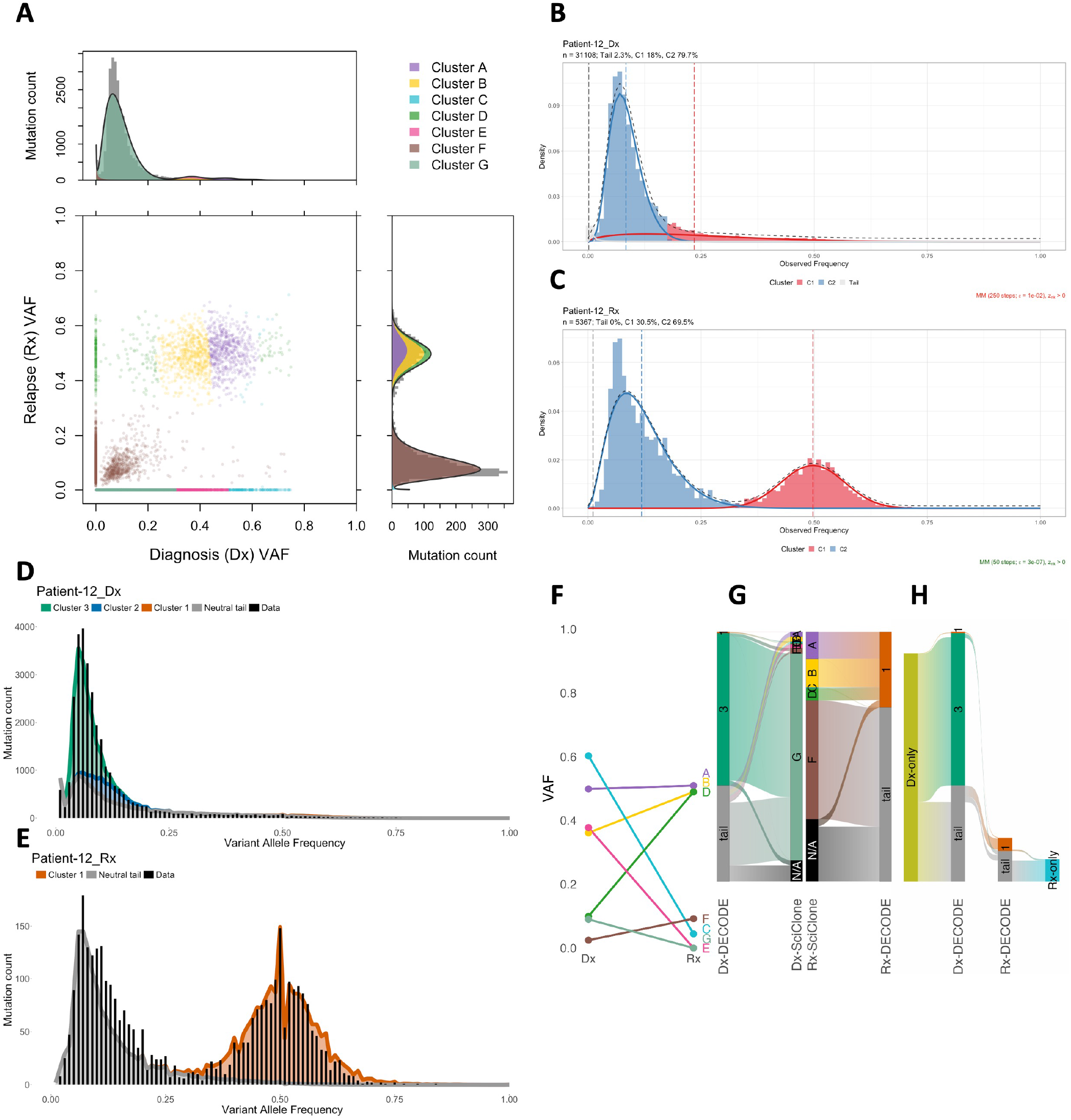
Mutation clustering for patient AML-12. **A**: SciClone’s joint clustering of mutations at diagnosis and relapse. **B-E**: Deconvolution of mutations using MOBSTER (**B, C**) and DECODE (**D, E**) for each sample at diagnosis (**B, D**) and relapse (**C, E**). **F**: Mean VAFs between diagnosis and relapse of subclones inferred from SciClone. **G**: Alluvial diagrams of subclonal assignments for mutations at diagnosis (left) and relapse (right) between DECODE and SciClone. Widths are proportional to mutation fractions. **H**: Alluvial diagram of DECODE subclonal assignments for mutations at diagnosis and relapse. Outer bars contain mutations unique to each sample.

**Supplementary Figure 15:**
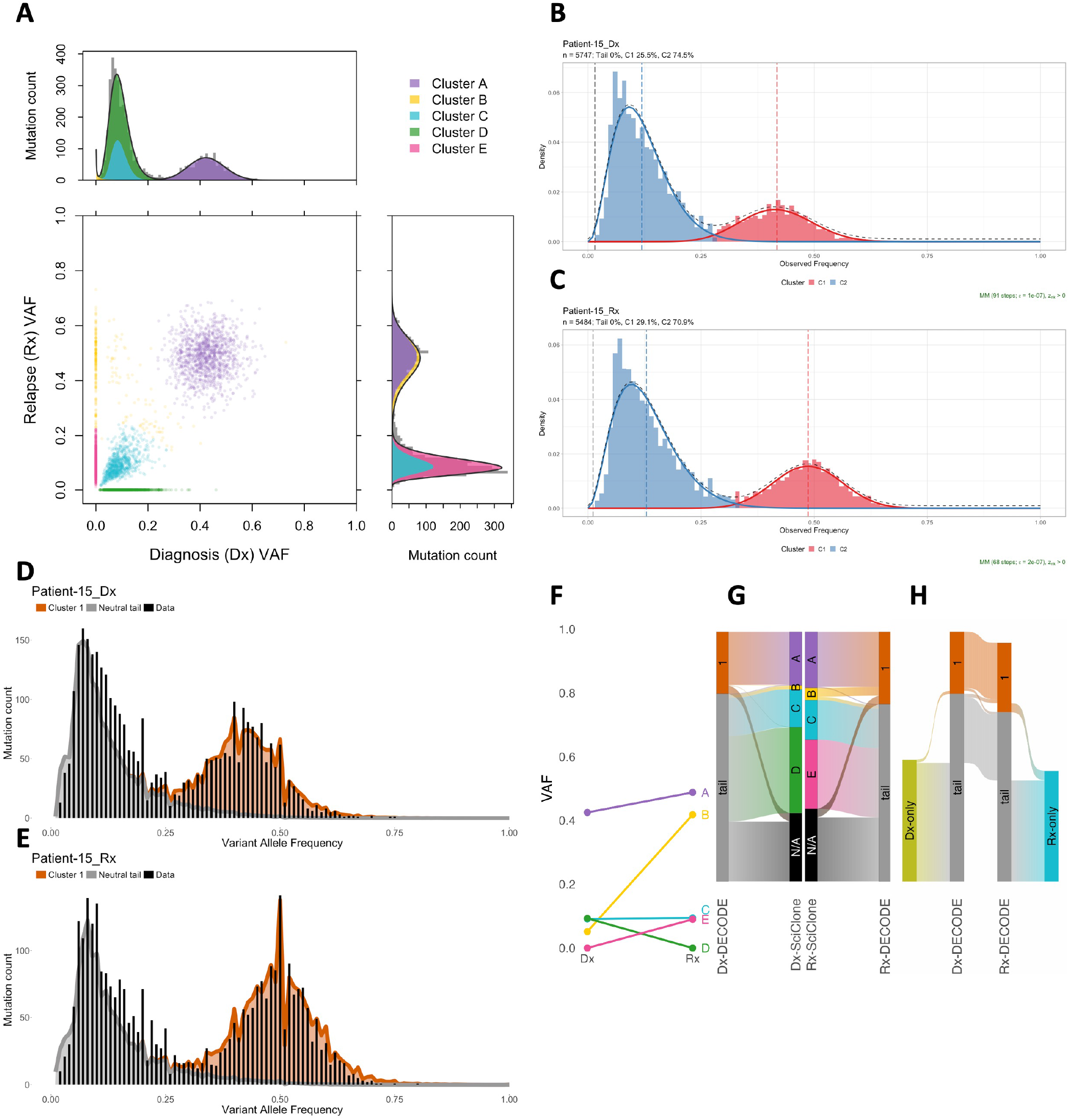
Mutation clustering for patient AML-15. **A**: SciClone’s joint clustering of mutations at diagnosis and relapse. **B-E**: Deconvolution of mutations using MOBSTER (**B, C**) and DECODE (**D, E**) for each sample at diagnosis (**B, D**) and relapse (**C, E**). **F**: Mean VAFs between diagnosis and relapse of subclones inferred from SciClone. **G**: Alluvial diagrams of subclonal assignments for mutations at diagnosis (left) and relapse (right) between DECODE and SciClone. Widths are proportional to mutation fractions. **H**: Alluvial diagram of DECODE subclonal assignments for mutations at diagnosis and relapse. Outer bars contain mutations unique to each sample.

